# High-throughput reclassification of *SCN5A* variants

**DOI:** 10.1101/858175

**Authors:** Andrew M. Glazer, Yuko Wada, Bian Li, Ayesha Muhammad, Olivia R. Kalash, Matthew J. O’Neill, Tiffany Shields, Lynn Hall, Laura Short, Marcia A. Blair, Brett M. Kroncke, John A. Capra, Dan M. Roden

## Abstract

**Rationale:** Partial or complete loss of function variants in *SCN5A* are the most common genetic cause of the arrhythmia disorder Brugada Syndrome (BrS1). However, the pathogenicity of *SCN5A* variants is often unknown or disputed; 80% of the 1,390 *SCN5A* missense variants observed in at least one individual to date are variants of uncertain significance (VUS). The designation of VUS is a barrier to the use of sequence data in clinical care.

**Objective:** We selected 83 variants for study: 10 previously studied control variants, 10 suspected benign variants, and 63 suspected Brugada Syndrome-associated variants, selected on the basis of their frequency in the general population and in patients with Brugada Syndrome. We used high-throughput automated patch clamping to study the function of the 83 variants, with the goal of reclassifying variants with functional data.

**Methods and Results:** Ten previously studied variants had functional properties concordant with published manual patch clamp data. All 10 suspected benign variants had wildtype-like function. 22 suspected BrS variants had loss of channel function (<10% normalized peak current) and 23 variants had partial loss of function (10-50% normalized peak current). The 73 previously unstudied variants were initially classified as likely benign (n=2), likely pathogenic (n=11), or VUS (n=60). After the patch clamp studies, 16 variants were benign/likely benign, 47 were pathogenic/likely pathogenic, and only 10 were still VUS. 8/22 loss of function variants were partially rescuable by incubation at lower temperature or pretreatment with a sodium channel blocker. Structural modeling identified likely mechanisms for loss of function including altered thermostability, and disruptions to alpha helices, disulfide bonds, or the permeation pore.

**Conclusions:** High-throughput automated patch clamping enabled the reclassification of the majority of tested VUS’s in *SCN5A*.

## Introduction

Variants in the cardiac sodium channel gene *SCN5A* (which encodes the protein NaV1.5) have been linked to many arrhythmia and cardiac conditions, including Brugada Syndrome (BrS),^1^ Long QT Syndrome (LQT),^2^ dilated cardiomyopathy,^3^ cardiac conduction disease,^4^ and Sick Sinus Syndrome.^5^ BrS is diagnosed by a characteristic ECG pattern (ST-segment elevation in the right precordial leads) at baseline or upon drug challenge, and is associated with an increased risk for sudden cardiac death.^6^ *SCN5A* loss of function variants are the most common (and in fact, the only ClinGen-validated) genetic cause of BrS, and are present in approximately 30% of cases (BrS1).^7, 8^ Variants that destabilize fast inactivation of the channel and generate a persistent (“late”) current are associated with LQT.^9, 10^ The risk of sudden cardiac death in BrS patients can be partially prevented, for example with implantable defibrillators. Therefore, incidentally discovered *SCN5A* variants that are pathogenic or likely pathogenic for BrS are considered actionable and reportable.^11^

A multi-center study of 2,111 unrelated individuals with Brugada Syndrome discovered 293 distinct *SCN5A* variants in these patients, including 225 variants present in only one patient.^1^ This result indicates that Brugada Syndrome risk is distributed among many different, rare *SCN5A* variants. In a recent curation of *SCN5A* variants, we identified 1,712 variants that have been observed to date in at least one individual in the literature or in the genome aggregation database (gnomAD).^9^ The strongest predictor of a variant’s association with Brugada Syndrome was a partial or total reduction in peak current recorded during voltage clamp experiments in heterologous expression systems (mainly HEK293 cells).^9^ Additionally, a weaker association was observed between a variant’s Brugada Syndrome association and its V_½_ activation, the voltage that elicits half-maximal channel activation.

Because *SCN5A* variants linked to Brugada Syndrome have incomplete penetrance, assessment of variant risk using phenotypic data in single variant carriers can be minimally informative.^12^ Nevertheless, an important challenge is to accurately classify variants, especially when only limited patient data are available.^13^ The American College of Medical Genetics and Genomics (ACMG) classification scheme uses 28 criteria to classify variants into 5 categories: pathogenic, likely pathogenic, benign, likely benign, or variant of uncertain significance (VUS).^14^ The guidelines have four “strong evidence” criteria towards classifying a variant as pathogenic: PS1 (same amino acid change as an established variant), PS2 (*de novo* in a patient with disease and no family history), PS3 (well-established functional variant studies show a deleterious effect), and PS4 (prevalence in cases statistically increased over controls). *In vitro* functional studies that demonstrate that a variant does not affect protein function can meet benign evidence criterion 3 (BS3, well-established functional variant studies show no deleterious effect). Thus, *in vitro* functional studies are a key method to aid in the classification of the large number of *SCN5A* variants.

Conventionally, variants in ion channels such as *SCN5A* have been assessed by time-intensive manual patch clamping of variants in heterologous expression. Recently, automated high-throughput patch clamp devices have allowed the rapid evaluation of dozens of variants in ion channel genes, including the arrhythmia-associated genes *KCNQ1* and *KCNH2*.^15-17^ In this study, we use automated patch clamping to study 83 *SCN5A* variants, identify 45 new partial or total loss of function variants, and reclassify 50/60 variants of uncertain significance.

## Methods

### Selection of variants to study

83 variants were studied by automated patch clamping (Figure 1): 10 previously studied controls, 10 suspected benign variants, and 63 suspected BrS variants. The previously studied variants were chosen because they had a spectrum of normal and abnormal channel function, representing multiple types of perturbations to the channel. The suspected BrS variants were selected because they appeared in at least 1 case of BrS in the literature and were relatively rare or absent in unaffected individuals in the literature and in the gnomAD database of putative controls (Figure 2B, Table S1).^9, 18^ In contrast, the suspected benign variants were observed in multiple unaffected or gnomAD individuals, but never in cases of BrS or LQT in the literature (Figure 2B, Table S1). R1958X was also classified as a suspected BrS variant because it is a nonsense variant. However, to our knowledge, it has not been observed in any patients with BrS (Figure 2B, Table S1).

**Figure 1.**
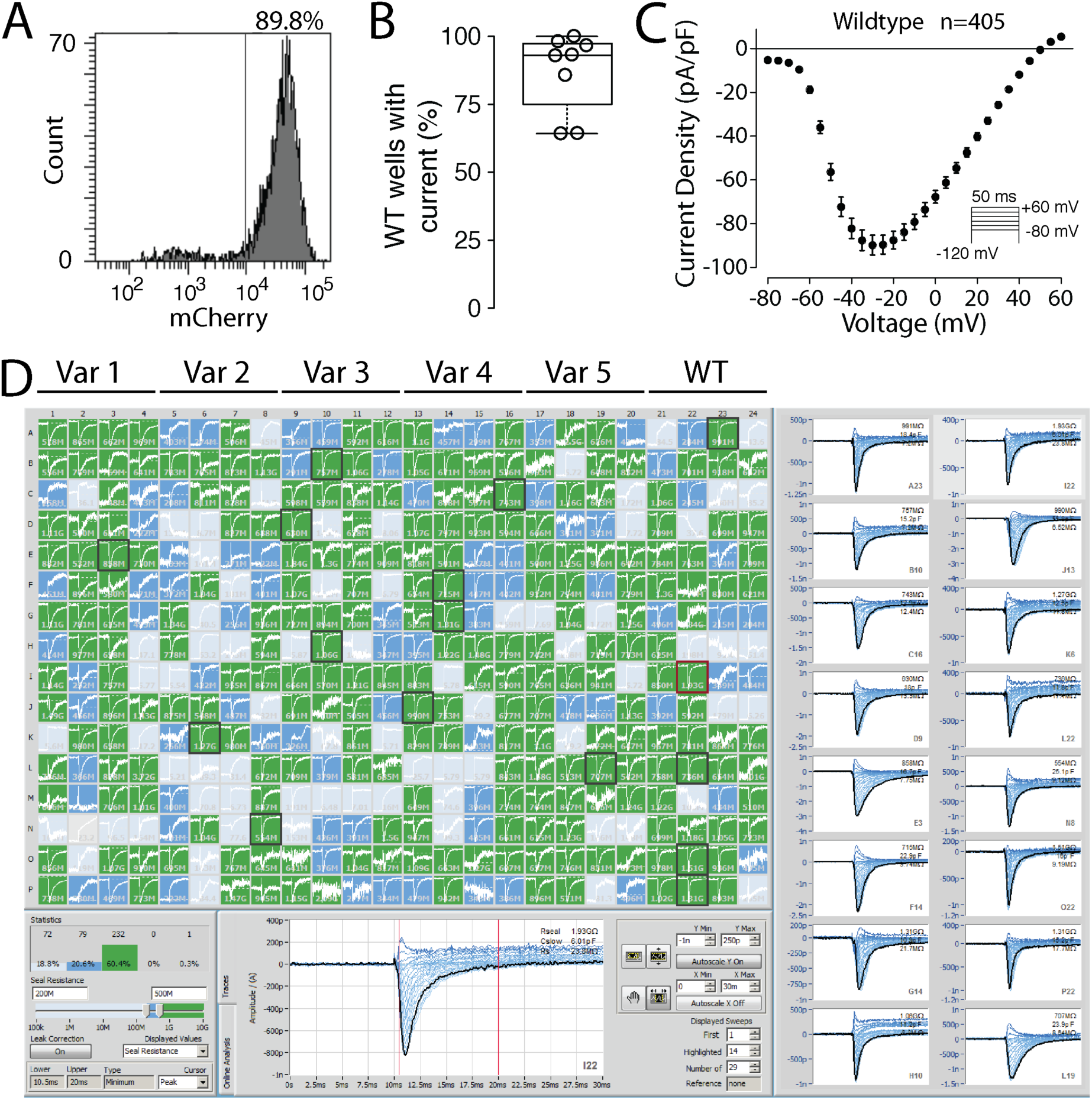
Molecular and functional expression of *SCN5A*. A) Flow cytometry plot of representative wildtype stable cell line. Histogram of mCherry signal is shown. 89.8% of cells expressed a high-level of mCherry, indicating successful plasmid integration. B) Boxplot of percentage of wildtype wells with NaV1.5-like current across 8 independent transfections. Only wells with a cell passing seal resistance and capacitance criteria (see Methods) were considered. C) Wildtype current density-voltage plot, averaged across 405 wildtype cells passing quality control (seal resistance, capacitance, in voltage control, and minimum peak current criteria, see Methods). The predicted reversal potential given the internal and external solutions used in this study is 45.3 mV. Error bars indicate s.e.m. D) Example SyncroPatch experiment. A typical experiment studied 5 variants and wildtype on a single 384-well chip. In this experiment all 5 variants had wildtype-like currents. Green wells indicate a cell with seal resistance >0.5 GΩ, blue wells indicate a cell with seal resistance 0.2-0.5 GΩ (not included in analysis), and grey wells indicate no cell present or a cell with seal resistance <0.2 GΩ (not included in analysis). Randomly selected cells with seals > 0.5GΩ are highlighted with a black square and displayed on the right.

**Figure 2:**
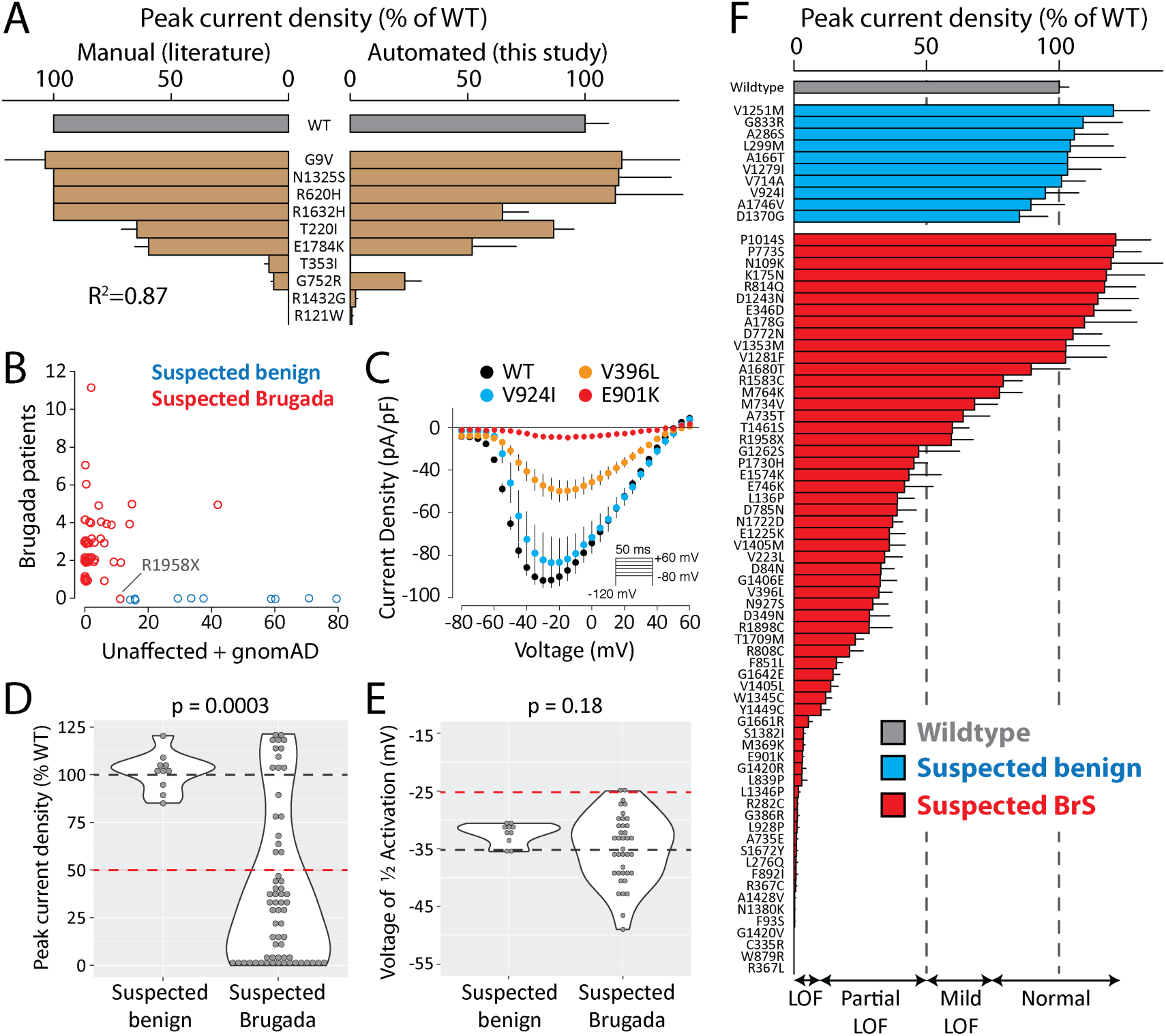
Suspected Brugada Syndrome variants have reduced peak current. A) Peak current density of 10 variants previously studied by manual patch clamp. Mean ± standard errors. Left: literature values. For some variants standard errors were not reported. Right: Automated patch clamp values (this study). B) Selection of suspected benign and suspected BrS-associated variants. Each data point is a variant. Points were “jittered” in the x- and y-axes with a small random number so that points with identical values would be visible. The y-axis indicates the number of patients with Brugada Syndrome in the literature.^9^ The x-axis is the number of unaffected individuals from the literature added to putative controls from the gnomAD database. Values are taken from a recent collation of *SCN5A* data from the literature.^9^ R1958X was classified as a suspected BrS variant because it is a nonsense variant, even though it has not been observed in any cases of BrS. C) Current-voltage curves for selected variants, showing an example of a normal variant (V924I), a partial loss of function variant (V396L), and a loss of function variant (E901K). D-E) Violin plots comparing peak current density (D) or voltage of ½ activation (E) for suspected Brugada Syndrome vs. suspected benign variants. Wildtype values are indicated with a dashed black line, and cutoffs for deleteriousness are indicated with a red line (50% peak current or 10 mV rightward shift in V_½_ activation). P values are from a two-tailed Mann-Whitney U test. For a complete list of measured parameters see Table S5. F) Peak current density (normalized to wildtype) for all previously unstudied variants. Error bar indicates standard error of the mean.

### *SCN5A* mutagenesis

To facilitate mutagenesis, a “zone” system was developed that divided the 6 kilobase *SCN5A* cDNA into 8 zones of ∼750 bp each (Table S2). These zones were separated by restriction enzyme sites already present in the *SCN5A* coding sequence or introduced with synonymous mutations. Mutagenesis primers were designed with the QuikChange primer design tool (https://www.agilent.com/store/primerDesignProgram.jsp). All mutagenesis primers are listed in Table S3. The QuikChange Lighting Multi kit (Agilent) was used to perform the mutagenesis with one primer/mutation, following manufacturer’s instructions. Mutants were created in a small (<4 kilobase) plasmid containing the zone of interest. Following sequence verification, mutant zones were moved with the appropriate restriction enzymes into a plasmid containing an AttB recombination site and a full length *SCN5A*:IRES:mCherry-blasticidinR (Figure S1). This plasmid was modified from a previously published promoterless dsRed-Express-derivative plasmid designed for rapid stable line generation.^19^ This modification introduced *SCN5A* into the plasmid and included the generation of a mCherry-blasticidinR fusion protein. For all plasmids, the entire *SCN5A* coding region was Sanger sequenced to confirm that the variant of interest was included and to verify that there were no off-target variants. The “wildtype” *SCN5A* used in this study had the more common H allele of the H558R polymorphism (rs1805124) and the most common splice isoform in the heart, a 2015 amino acid sequence (not counting the stop codon) that includes deletion of the Q1077 residue (ENST00000443581). However, the variants are named according to the 2016 amino acid Q1077-included isoform (ENST00000333535), as is standard in the literature.

### Stable *SCN5A*-expressing cell lines

Human Embryonic Kidney HEK293T “negative selection” landing pad cells were used, which express Blue Fluorescent Protein and an activatable iCasp caspase from a tetracycline-sensitive promoter.^20^ In these cells, an AttP integration site is located in between the promoter and the BFP/iCasp. Bxb1 integrase-mediated integration of a plasmid containing an AttB site at the genomic AttP location disrupts the expression of BFP/iCasp. After integration of an AttB-containing *SCN5A*:IRES:mCherry:blasticidinR plasmid, these cells switch to expression of *SCN5A* and an mCherry-blasticidinR fusion protein. Cells were cultured at 37°C in a humidified 95% air/5% CO2 incubator in “HEK media”: Dulbecco’s Eagle’s medium supplemented with 10% fetal bovine serum, 1% non-essential amino acids, and 1% penicillin/streptomycin. On day -1, cells were plated in 12 well plates. On day 0, each well was transfected with 2.1 ul FuGENE 6 (Promega), 600 ng of the mutant *SCN5A* plasmid and 100 ng of a plasmid expressing Bxb1 recombinase (a gift from Pawel Pelczar,^21^ Addgene plasmid #51271). On day 1, cells were washed with HEK media to remove the FuGENE reagent. On day 3, the cells were exposed to 1 μg/ml doxycycline (Sigma) in HEK media to induce expression of either the *SCN5A* mutant or iCasp9 in non-integrated cells from the Tet-sensitive promoter. On day 5, cells were exposed to 1 μg/ml doxycycline, 100 μg/ml blasticidin S (Sigma), and 10 nM AP1903 (MedChemExpress) in HEK media. The blasticidin selected for expression of the blasticidin resistance gene present on the *SCN5A* plasmid. The AP1903 activated the iCasp caspase protein expressed from non-integrated cells, this resulted in selection against non-integrated cells. Cells were exposed to doxycycline+blasticidin+AP1903 in HEK media for 4-7 days to select for integrated cells, passaging cells as needed. Less than 48 hours prior to the SyncroPatch experiment, cells were analyzed by analytical flow cytometry on a BD Fortessa 4-laser instrument to determine the percentage of cells that were mCherry-positive, Blue Fluorescent Protein-negative, indicating cells with successful integration of the *SCN5A* plasmid. BFP was measured by excitation at 405 and emission at 450/50 and mCherry was measured by excitation at 561 and emission at 610/20.

### Automated patch clamping

Cells were patch clamped with the SyncroPatch 384PE automated patch clamping device (Nanion). To prepare cells for patch clamping, cells were washed in PBS, treated with Accutase (Millipore-Sigma) for 3 minutes at 37°C, then recovered in CHO-S-serum free media (Gibco). Cells were pelleted and resuspended in divalent free reference solution (DVF) at ∼200,000-400,000 cells/ml. DVF contained (mM) NaCl 140, KCl 4, alpha-D(+)-glucose 5, HEPES 10, pH 7.4 adjusted with NaOH. Cells were then added to a medium resistance (4-6 MΩ) 384-well recording chamber with 1 patch aperture per well (NPC-384, Nanion), which contained DVF and internal solution: CsCl 10, NaCl 10, CsF 110, EGTA 10, HEPES 10, pH 7.2 adjusted with CsOH. Next, to enhance seal resistance, 50% of the DVF was exchanged with a calcium-containing seal enhancing solution: NaCl 80, NMDG 60, KCl 4, MgCl2 1, CaCl2 10, alpha-D(+)-glucose 5, HEPES 10, pH 7.4 adjusted with HCl. The cells were washed three times in external recording solution: NaCl 80, NMDG 60, KCl 4, MgCl2 1, CaCl2 2, alpha-D(+)-glucose 5, HEPES 10, pH 7.4 adjusted with HCl. Currents elicited in response to activation, inactivation, and recovery from inactivation protocols were then recorded (Figure S2). A late current measurement was captured every 5 seconds. After 1 minute, 50% of the external solution was exchanged with external solution containing 200 μM tetracaine hydrochloride (Sigma; effective concentration 100 μM tetracaine). After tetracaine addition, late current measurements were obtained every 5 seconds for 1 additional minute. At least 10 cells expressing wildtype *SCN5A* were included for comparison in each SyncroPatch experiment (Figure 1), and at least 2 independent transfections and at least 10 replicate cells were studied per mutant. Recordings were performed at room temperature.

We also conducted experiments to assess the effects of incubation at low temperature or mexiletine (a sodium channel blocker), interventions reported to increase cell surface expression of mistrafficked channels.^22-26^ For these experiments, cells stably expressing loss of function variants were generated as described above. The cells were incubated for 24 hours at 30°, or at 37° with or without 500 μM mexiletine hydrochloride (Sigma), washed with HEK media, and were patch clamped as described above.

### Data analysis

Cells with a seal resistance of 0.5-10 GΩ and a cell capacitance between 5-30 pF were analyzed. Built-in SyncroPatch windowing methods were used to calculate currents in peak/inactivation/recovery from inactivation protocols (maximum current from 0.5-5 ms post voltage shift) or late current (mean current from 190-200 ms). These values were exported to .csv files and analyzed in custom R scripts. Current-voltage plots were created for each well, and each well was manually analyzed and classified as normal/in voltage control (voltage-dependent current with peak current near -20 mV), out of voltage control (voltage-dependent current that rapidly jumped around -60 mV), or having no current (<25 pA peak current). Example current-voltage curves of these three classes are shown in Figure S3.

Only wells that were in voltage control (Figure S3) with peak currents between 100-2000 pA were used to assess additional features, such as the voltage dependence of activation, voltage dependence of inactivation, inactivation time, or recovery from inactivation. Additional details on peak current averaging are presented in the Supplemental Methods. Only wells with a peak current above 1000 pA were used to measure late current. Activation and inactivation best-fit curves were calculated for each well by fitting Boltzmann equations using the R function *nls* (nonlinear least squares). Recovery from inactivation data and inactivation time data were fitted with exponential curves with the *nls* function. Wells with high noise for which the best-fit curve did not fit the data well (data points had >10% average deviation from the best-fit line) were removed from the analysis. Tetracaine-sensitive late current was calculated as the mean of 5 post-tetracaine raw late current values subtracted from the mean of 5 pre-tetracaine values and was normalized to tetracaine-sensitive peak current. Outlier values exceeding 3 standard deviations from the mean were excluded (0.9% of all values). Per-well V_½_ activation (voltage at which half the channels are activated), V_½_ inactivation (voltage at which half the channels are inactivated), time of 50% recovery from inactivation, inactivation time constant, and late current parameters were averaged across all wells meeting the above inclusion criteria. All measured parameters with at least 5 cells meeting inclusion criteria are reported. For many severe loss of function variants there were fewer than 5 qualifying wells to accurately quantify channel parameters other than peak current density. All comparisons of variant parameters between groups were made in R with two-tailed *t*-tests (*t*.*test*), or with two-tailed Mann-Whitney U tests (*wilcox*.*test* with the parameter paired=FALSE) when the distributions were non-normally distributed. Differences in dispersion between groups were tested with Levene’s Test (*levene*.*test, car* package). Violin plots were made with *geom_violin* (ggplot2).

Variants were classified according to American College of Medical Genetics and Genomics (ACMG) criteria (Figure S4).^14^ A custom R script was used to implement these criteria. Variant classifications pre- and post-functional data are presented in Table S1. A cutoff of 6/∼250,000 alleles in gnomAD v.2.1^18^ was used to determine criteria BS1 and PM2, following a previous recommended cutoff for Brugada Syndrome.^27^ BP4 and PP3 were determined from the consensus of PROVEAN^28^ and PolyPhen2^29^ classifications. PS4 was interpreted to mean at least 5 observed individuals with Brugada Syndrome and an estimated Brugada Syndrome penetrance from literature reports whose 95% confidence interval excluded 0.^9^ Variants with peak current densities between 75-125% of wildtype, <10 mV shifts in activation or inactivation, <2-fold shifts in recovery from inactivation, and <1% late current (% of peak) were considered to have normal *in vitro* functional data (BS3). Variants with peak current densities <50% of wildtype or a >10 mV rightward shift in V_1/2_ activation were considered to have abnormal loss of function functional data (PS3). Variants with >75% peak current and a late current >1% (normalized to peak current) were considered to have abnormal gain of function functional data (PS3). These cutoffs were determined from a previous analysis of the correlation between functional parameters and Brugada Syndrome and Long QT Syndrome risk.^9^ ClinVar classifications^30^ were used to determine criteria BP6 and PP5. All literature and gnomAD patient counts, peak current densities, and classifications pre- and post-patch clamp data are presented in Table S1.

### Homology model and structural calculations

All computational modeling was conducted in parallel to and blinded from the experimental characterizations. Structural models of human *SCN5A* (UniProtKB accession number: Q14524-1, modeled residues: 30-440, 685-957, 1174-1887) bound with *SCN1B* (UniProtKB accession number: Q07699-1, modeled residues: 20-192) were generated by homology modeling using the protein structure prediction software Rosetta (version 3.10).^31^ The cryo-EM structure of human *SCN9A* bound with *SCN1B* and the Ig domain of *SCN2B* resolved to 3.2 Å (PDB 6J8H)^32^ was used as the primary template while the cryo-EM structure of NavPaS from American Cockroach resolved to 2.6 Å (PDB ID: 6A95)^33^ was used as a secondary template. The percent identity between the aligned positions of *SCN9A* and *SCN5A* sequences was 76.7%. While the percent identity between NavPaS and *SCN5A* was only moderate (45.6%), the N-terminal and C-terminal domains in the NavPaS structure were partially resolved, providing coordinates for modeling the corresponding domains of *SCN5A*. The final model (Figure 5, S5) covers 70 of the 73 functionally characterized variants. Three variants, P1014S, R1898C, and R1958X fall outside of the set of modeled residues, and are therefore not covered. Additional information about the homology model and ΔΔG (thermostability) calculations are presented in the Supplemental Methods and Table S4.

**Figure 3:**
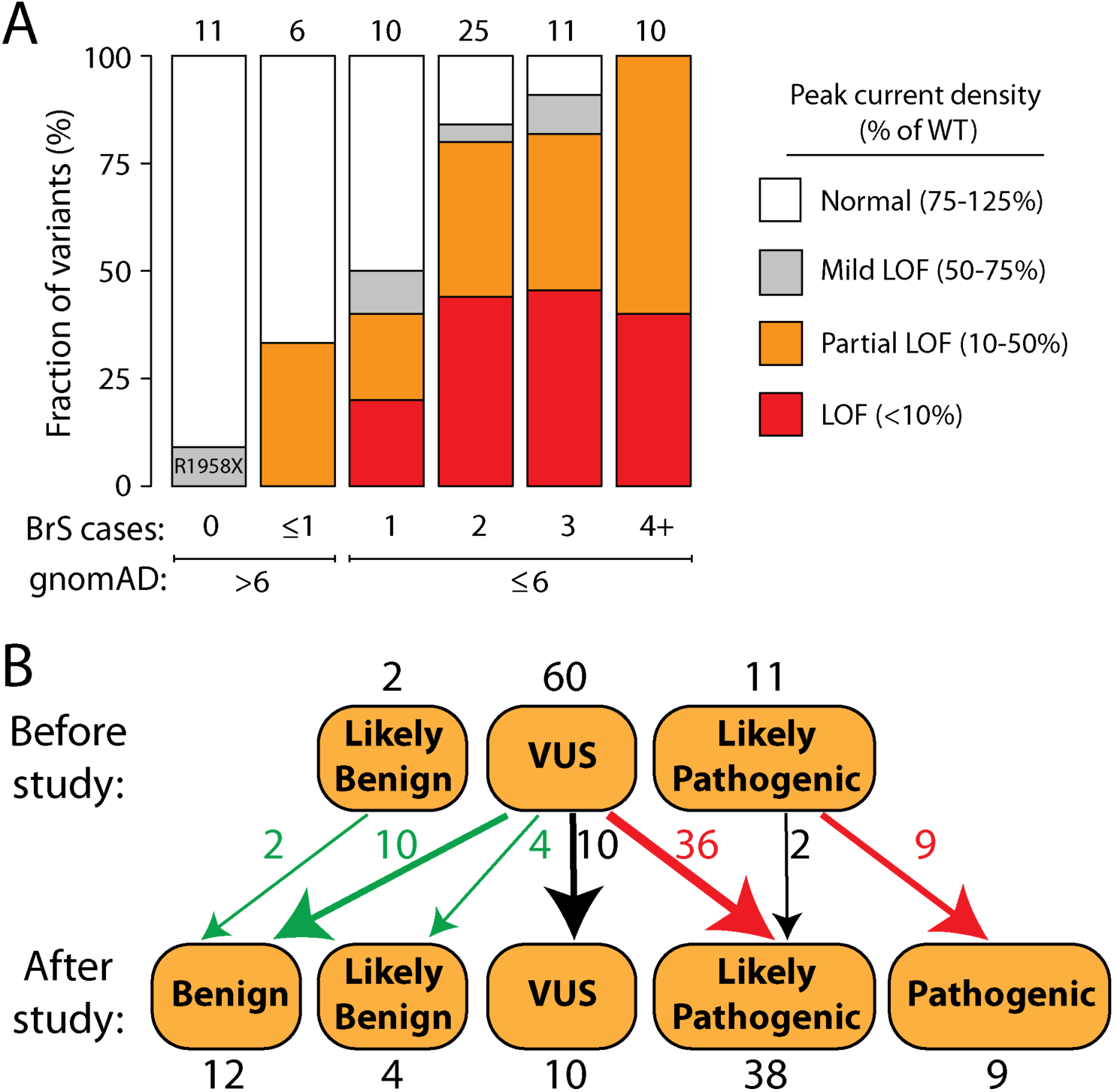
Reclassifications with patch clamp data. A) Patch clamp results varied depending on the number of observed Brugada patients in the literature^9, 13^ and the gnomAD allele frequency.^18^ LOF=loss of function. B) Variants were classified according to the American College of Medical Genetics and Genomics criteria.^14^ Above: classifications before the patch clamp data in this study. Below: classifications with the patch clamp data in this study. All classifications indicate Brugada Syndrome, except for R814Q, which was reclassified from Likely Pathogenic for Brugada Syndrome to Likely Pathogenic for Long QT Syndrome.

**Figure 4.**
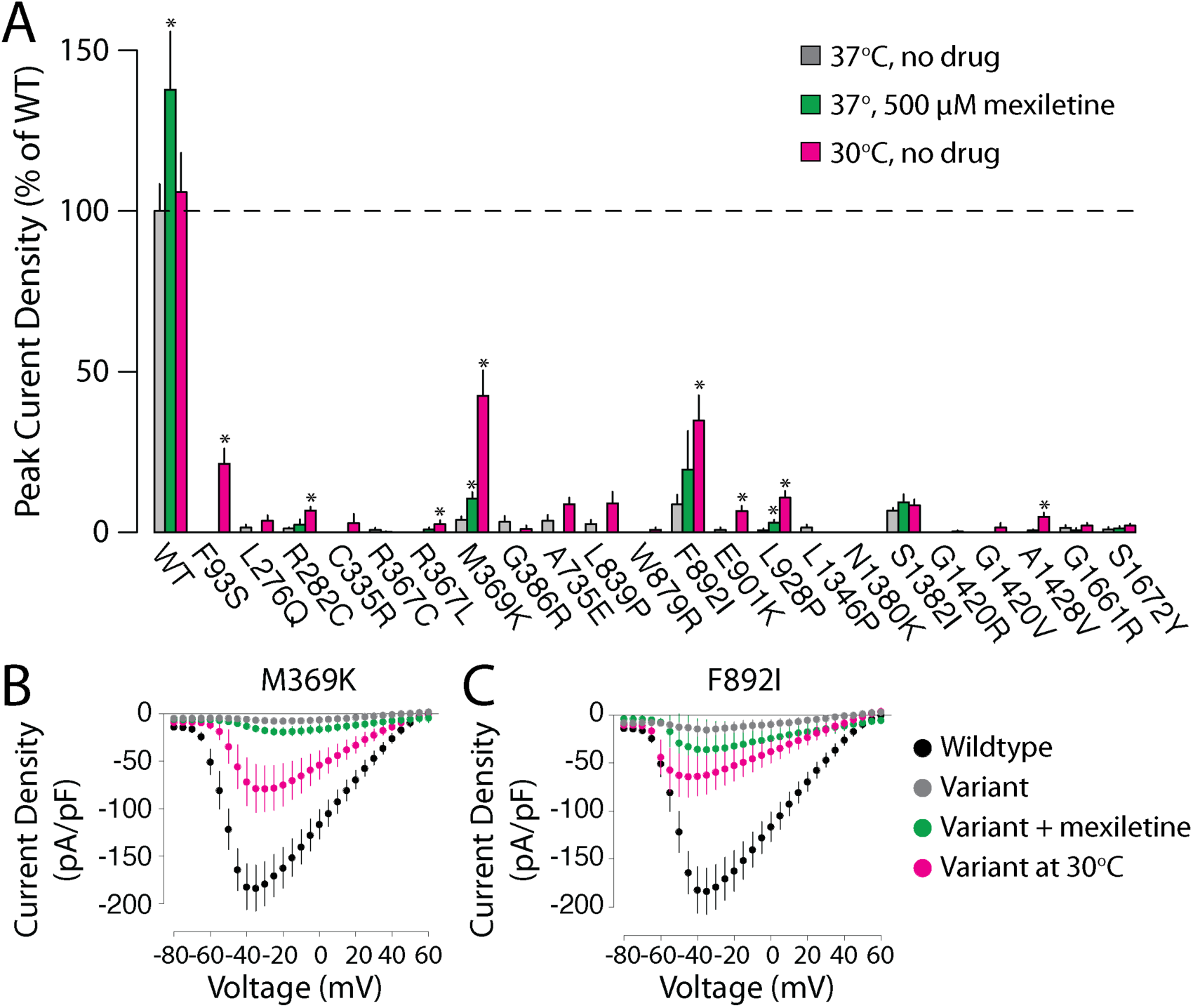
Partial rescue of loss of function variants. A) Peak current densities (normalized to wildtype-expressing cells from the same transfection) for the 22 variants with peak current densities <10% of wildtype. Variants were tested under usual conditions (37°, no drug, grey), with 24 hr of treatment with 500 μM mexiletine (green), or with 24 hr culture at 30° (pink). Values are mean ± standard error. *P<0.05, 2-tailed *t*-test compared with wildtype at 37° with no drug. 8/22 variants had significant increases in current at 30° and 2/22 variants had significant increases in mexiletine. B, C) Current-voltage plots for M369K and F892I, the two variants with the largest response to 30°. Colors are the same as A), with black indicating wildtype at 37° with no drug.

**Figure 5.**
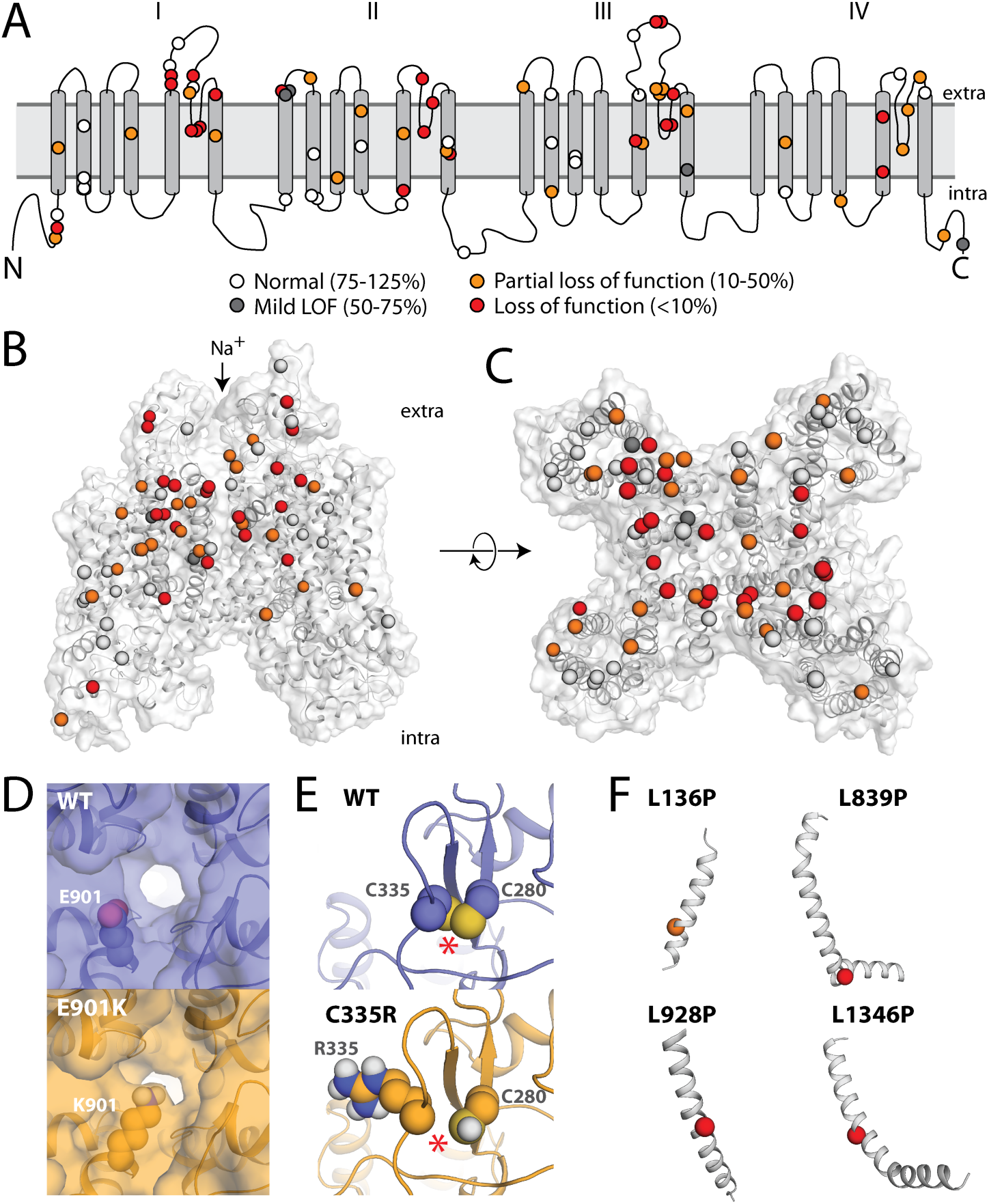
Structural basis of *SCN5A* loss of function variants. A) Two-dimensional schematic of NaV1.5 structure. All previously unstudied variants are shown and color-coded based on peak current density (white 75-125%, grey 50-75%, orange 10-50%, red <10%). B, C) Three-dimensional homology model of NaV1.5. Variants are colored as in A. D) Top-down view of WT (top) and E901K (bottom), as modeled using Rosetta. The lysine residue projects into the pore, likely disrupting sodium passage. E) View of WT and C335R, as modeled using Rosetta. The WT protein has a disulfide bond between C335 (left) and C280 (right), which was inferred from the spatial proximity of these two residues and the fact that the corresponding residues in the template structures also form a disulfide bond; this bond is disrupted by C335R. The disulfide bond is indicated with *. F) Four leucine->proline variants in this study. L136P is a partial loss of function variant and L839P, L928P, and L1340P are loss of function. The structures of these four variants were not modeled because modeling drastic structural changes involving prolines that are part of a helix usually cannot be reliably modeled in Rosetta. However these variants likely cause loss of protein function by causing a kink in the alpha helix and protein misfolding.

## Results

### Automated patch clamping of NaV1.5 variants

Across multiple transfections and variants, a high percentage (85.4±12.3% s.e.m.) of cells were mCherry-positive, indicating successful integration of an *SCN5A* expression plasmid (Figure 1A, S1). Wildtype and mutant cells were patch clamped with the SyncroPatch 384PE device (Figure 1). For cells expressing wildtype *SCN5A*, most wells that passed seal resistance and capacitance quality control cutoffs had voltage-dependent sodium currents (86.9% ± 14.6% s.e.m., Figure 1B-C). Wildtype cells had typical NaV1.5 properties (Figure 1, S2).

To validate the automated patch clamping method, we patch clamped 10 previously studied variants chosen because they had a range of functional properties. The previously published manual patch clamp properties were concordant with the automated patch clamp data (Figure 2A, Table S5). Peak current is the major parameter of interest for assessing Brugada Syndrome risk.^9^ We observed a strong correlation (R^2^=0.87) between the automated peak current density measurements and previously published manual patch clamp peak current density values (Figure 2A).

From our previous curation of the *SCN5A* variant literature,^9^ we selected 73 previously unstudied variants: 63 suspected Brugada Syndrome variants and 10 suspected benign variants. The suspected benign and suspected Brugada Syndrome variants were selected based on their frequency in the gnomAD database and in the literature in patients with Brugada Syndrome. (Figure 2B, Table S1).^9, 18^ Applying ACMG classification criteria, without patch clamping data, 60 variants were classified as VUS, 2 as likely benign, and 11 as likely pathogenic (Figure 3, Table S1).

### Most suspected Brugada Syndrome variants had (partial) loss of function

There was a striking difference in peak current density between the suspected BrS and suspected benign variants (p=0.0003, two-tailed Mann-Whitney U test, Figure 2D). Overall, 23/63 suspected BrS variants had partial loss of function (10-50% peak current density, normalized to wildtype) and 22/63 had loss of function (<10%), while 10/10 suspected benign variants had normal peak currents (75-125%) (Figure 2F, Table 1). We next examined whether large changes to other parameters were a common cause of channel loss of function. There was a statistically significant increase in width of the distribution of V_½_ activations in the suspected BrS set (p=0.0028, Levene’s test). However, only 2/39 variants with measured V_½_ activation had a >10 mV rightward shift (WT: 36.7±0.6 mV, n=279; D785N: -26.4±1.3 mV, n=16; D349N: -26.5±3.1 mV, n=5). D785N and D349N both also had a partial loss of function in peak current (38.9% and 28.5% of WT, respectively; Figure 2F). Overall, there was no significant difference in V_½_ activation between suspected BrS and suspected benign variants (p=0.18, Mann-Whitney U test, Figure 2E). Therefore, in this dataset, large shifts in activation gating were not a common cause of Brugada Syndrome. Surprisingly, one suspected BrS-associated variant, R814Q, had near wild-type-like peak current density (117% of wildtype) and increased late current (1.4% of peak). In addition to Brugada Syndrome cases, R814Q has been observed in two cases of Long QT Syndrome,^34^ consistent with its gain of function late current phenotype. Besides these variants, no major differences between suspected BrS and suspected benign variants were observed for V_½_ inactivation, inactivation time, recovery from inactivation, or late current (Figure S6-10, Table S5). Despite being a nonsense variant, R1958X generated substantial current (peak current density of 59.3% of wildtype), likely due to the fact that the stop codon is in the distal C-terminus of the protein (Figure 2).

**Table 1:** Functional class of suspected benign and Brugada Syndrome variants.

| **Functional class** | **Peak current density** | **Suspected benign** | **Suspected BrS** |
| --- | --- | --- | --- |
| Normal | 75-125% | 10 (100%) | 14 (22%) |
| Mild loss of function | 50-75% | 0 (0%) | 4 (6%) |
| Partial loss of function | 10-50% | 0 (0%) | 23 (37%) |
| Loss of function | <10% | 0 (0%) | 22 (35%) |
| Total | * | 10 | 63 |

Some suspected BrS variants have only been observed in only a single BrS patient, whereas other variants have been observed in multiple individuals. In this study, the patch clamp phenotypes varied according to the strength of the phenotypic evidence for Brugada Syndrome (Figure 3A). For example, 4/10 variants that have been observed in exactly 1 case of Brugada Syndrome and ≤6 in gnomAD had partial or complete loss of function (0-50% peak current). In contrast, 10/10 variants that have been observed in at least 4 BrS1 cases and ≤6 in gnomAD had partial or complete loss of function. 2/6 variants seen in ≥ 1 cases of Brugada Syndrome but also in >6 counts in gnomAD had partial loss of function defects. Therefore, variants that were more commonly observed in Brugada Syndrome cases and less frequently observed in the population were more likely to have loss of channel function.

### Reclassification of variants with functional data

When we implemented the ACMG classification criteria^14^ to classify all studied variants without the automated patch clamp data (Figure 3B), 60/73 variants were classified as VUS. Based on our previous analysis of the relationship between functional perturbations and Brugada Syndrome risk,^9^ we defined cutoffs for ACMG functional criteria PS3 and BS3. Variants with <50% peak current were considered to meet criterion PS3 (well-established functional assays show a change), and variants with 75-125% peak current were considered to meet criterion BS3 (well-established functional assays show no change). Because of its elevated late current and normal peak current, R814Q was considered to meet the PS3 criterion for abnormal gain of function, and was reclassified from likely pathogenic for Brugada Syndrome to likely pathogenic for Long QT Syndrome. After patch clamping data were incorporated, 36 VUS’s were reclassified as likely pathogenic and 14 were reclassified as likely benign or benign. Overall, classifications were changed for 62/73 (85%) previously unstudied variants. Therefore, for this set of variants, functional data led to reclassification of the great majority of VUS’s.

### Partial rescue of some loss of function variants

We also tested whether incubating cells at low temperature or with a sodium channel blocker could partially rescue sodium current for the 22 loss of function variants (<10% normalized peak current density). Cells expressing these variants were cultured for 24 hours in usual conditions at 37° with no added drug, at 30°, or with 500 μM mexiletine. Compared to usual conditions, 8/22 variants had significantly increased peak current at 30°, and 2/22 variants had significantly increased peak current when treated with mexiletine (Figure 4). F892I and M369K had the largest responses to 30° incubation, with normalized peak current density increasing from 8.7% at 37° to 34.8% at 30° for F892I (p=0.006, two tailed *t*-test) and from 3.9% to 42.5% for M369K (p=0.0003; Figure 4A-C).

### Structural basis of loss of function

In this study, 22 previously unstudied variants had <10% peak current, and an additional 23 variants had 10-50% current. We explored the structural basis of these variants’ decreased channel function. Although the suspected BrS variants were selected for study independent of their position in the protein, 42/45 (partial) loss of function variants were located in the four structured transmembrane domains, as opposed to unstructured linker regions or the N- and C-termini (Figure 5A-C). 33/45 (73%) loss/partial loss of function variants were located in the pore-forming or pore-adjacent S5, S5-S6 linker, or S6 regions, areas that we have previously identified as a hotspot for Brugada variants.^9, 12^ Indeed, variant distance from the pore in the protein structure was strongly correlated with normalized peak current density (Pearson’s r = 0.538, p = 1.33e-6, Figure S11).

Many disease-causing variants cause amino acid substitutions that lead to a significant perturbation to native thermostability (|ΔΔG|) of protein structure.^35, 36^ To investigate the extent to which the perturbation to the native thermostability of NaV1.5 is correlated with molecular function, we evaluated the impact of each variant on thermostability relative to the wild-type structure with a Rosetta ΔΔG protocol.^37^ Normalized peak currents and |ΔΔG| values were significantly negatively correlated (Pearson’s r = -0.31, p = 0.0092, Figure S12). The |ΔΔG| of variants with functional effects on peak current (peak current density <50%, n=44, median |ΔΔG|=2.00 kcal/mol) was significantly larger than normal/mild loss of function variants (peak current density >=50%, n=26, median |ΔΔG|=0.88 kcal/mol, Mann-Whitney U test, p=0.0031, Figure S12B). This result suggests that variant-induced disruption of native thermostability of NaV1.5 may be a major factor contributing to compromised function.

Variants can cause complete or partial loss of function through mechanisms other than affecting thermostability, such as disrupting the native topology or electrostatic environment of the pore. Using a homology model of NaV1.5, several variants had plausible mechanisms that could explain their loss of function phenotype (Figure 5D-F). Seven pore-lining residues in this study, D349N, R367C, R367L, W879R, R892I, E901K, and T1709M, caused either complete or partial loss of function while inducing negligible or only minor perturbations to native thermostability (Figure 5, S13). In particular, the residue E901 lines the channel pore, and in the homology model, the loss of function variant E901K projects into the pore, likely disrupting sodium permeation (Figure 5D). C335R disrupts a disulfide bond, likely destabilizing the tertiary structure of the protein and explaining its loss of function phenotype (Figure 5E). Three loss of function leucine→proline variants (L839P, L928P, and L1346P) and a fourth partial loss of function variant (L136P) likely disrupt alpha helices, as is seen in other proteins with proline variants (Figure 5F).^38^ These data indicate that structural features can help predict or explain channel loss of function in *SCN5A* variants.

## Discussion

### Reclassification of Brugada Syndrome variants with patch clamping data

This study identified 45 novel (partial) loss of function variants by high-throughput, automated patch clamping. This nearly doubles (from 24 to 46) the number of known loss of function missense *SCN5A* variants with <10% peak current (Table S6). As a result of the patch clamp data, 36 novel pathogenic/likely pathogenic variants and 14 novel benign/likely benign variants were identified. Overall, 62/73 variants, including 50/60 VUS’s, were reclassified with our patch clamp data. R1958X has been observed in 13 individuals in the gnomAD database and never in a published case of Brugada Syndrome. R1958X generated substantial current (peak current density of 59.3% of wildtype), and likely escapes Nonsense Mediated Decay because it located is in the last exon of *SCN5A* (Figure 2). Eight other nonsense and frameshift variants in the C-terminus of *SCN5A* are present in the gnomAD database;^18^ these variants may also generate sodium current and not have complete loss of channel function.

It is important to note that *SCN5A* variant classification does not completely predict Brugada Syndrome risk. BrS is an incompletely penetrant disease, and its presentation is influenced by demographic factors such as age and sex,^39^ as well as common genetic variants—including multiple noncoding haplotypes near *SCN5A*.^40^ Patients with loss of function *SCN5A* variants can present with other arrhythmias besides Brugada Syndrome, including Sick Sinus Syndrome,^5^ atrial standstill,^41^ or other conduction disease.^42^ Therefore, carriers of the loss of function variants identified in this study may present with those conditions instead of Brugada Syndrome. In addition, some “overlap” *SCN5A* variants can have increased risk of both Brugada Syndrome and Long QT Syndrome.^43^ One suspected BrS-associated variant in this study, R814Q, had late current above our cutoff (>1% of peak current), and has appeared in the literature in both BrS and LQT patients.^34, 44^ Patients with pathogenic or likely pathogenic *SCN5A* variants should have follow-up ECG screening and a clinical and family history taken to determine each individual’s phenotype and sudden cardiac death risk.

### Mechanisms of NaV1.5 loss of function

In this set of variants, the most common cause of channel loss of function was a partial or total reduction in peak current. This is consistent with our previous analysis which found that *SCN5A* peak current was the largest single predictor of Brugada Syndrome risk.9 Two variants, D785N and D349N, had a >10 mV rightward shift in V_½_ activation; however these two variants also had <50% peak current. Thus, it appears that while variants with large gating defects have been previously described (e.g. R1632H^45^), the major mechanism of NaV1.5 loss of function is a reduction in peak current. Previous studies have found that channel misfolding and a resulting cell-surface trafficking deficiency is a common mechanism of loss of function variants in *SCN5A*^5, 22^ and other ion channel genes.^46^ We observed a negative correlation between peak current density and |ΔΔG|, a parameter that indicates variant-induced perturbations to native thermostability, consistent with other studies that showed that thermostability perturbations are a major cause of altered protein function.^47, 48^ In addition, structural modeling identified several probable modes of channel dysfunction, including pore-lining variants that likely disrupt sodium permeation, removal of a disulfide bond, or creation of prolines that likely disrupt alpha helices. Although structural analyses were enlightening, these features do not yet fully predict channel function, indicating the necessity of empirical electrophysiological measurements. Consistent with previous studies of *SCN5A* variants,^22-26^ some but not all (8/22) *SCN5A* loss of function variants were partially rescued by 30° treatment or mexiletine. Although these treatments are not practical in the clinic for treatment of Brugada Syndrome, this result suggests that other drugs or interventions may help rescue some *SCN5A* loss of function variants.

### High-throughput approaches to variant classification

Variants in *SCN5A* account for approximately 30% of cases of Brugada Syndrome.^8^ The ClinGen consortium recently affirmed *SCN5A* as the only gene associated with Brugada Syndrome that meets preset criteria for disease association.^7^ A major challenge for the field is to identify which *SCN5A* variants are disease-associated. 1,712 *SCN5A* variants (including 1,390 missense variants) have been observed in at least one individual to date, most of which are VUS’s.^9^ The risk for Brugada Syndrome is spread over hundreds (and possibly thousands) of variants, with no single *SCN5A* variant present in >1% of cases.^1^ This study helps tackle the VUS problem for *SCN5A* by reclassifying 50/60 VUS’s, 36 to likely pathogenic and 14 to (likely) benign.

Previous studies have implemented a similar automated patch clamping approach on dozens of variants in two other arrhythmia-associated genes, *KCNQ1* and *KCNH2*.^16, 17^ Therefore, targeted high-throughput patch clamping of ion channel variants is a promising method for reclassifying additional variants in *SCN5A* and other Mendelian arrhythmia genes. This approach may be extended to study additional *SCN5A* variants, including gain of function Long QT-associated variants or additional variants observed in population sequencing efforts. Accurate classification of the large number of variants in arrhythmia-associated genes will require integrating data from multiple different model systems, such as patch clamping,^16, 17^ induced pluripotent stem cell-derived cardiomyocytes,^49-51^ structural and computational models,^12, 52, 53^ and ultra-high-throughput multiplexed assays.^54-56^

### Limitations

This study assays variants in a heterologous expression system. While peak current as measured in this system is the strongest available predictor of Brugada Syndrome risk,^9^ it is possible for some variants to show different properties in HEK293T cells compared to cardiomyocytes, e.g. because of other proteins that interact with or modify NaV1.5.57-59 This study only examined a single splice isoform and the most common haplotype (H558) of *SCN5A*; for some variants, it has been shown that alternate haplotypes/isoforms can affect channel properties.^60^

### Conclusion

This study used automated patch clamping to study 73 previously unstudied *SCN5A* variants, resulting in the reclassification of 50/60 variants of uncertain significance. This approach can help reclassify variants in this important disease gene and improve the accuracy and scope of genetic medicine.

## Supporting information

FileS1_EPdata

## Non-standard Abbreviations and Acronyms

ACMG: American College of Medical Genetics and Genomics;
BrS: Brugada Syndrome;
HEK: Human Embryonic Kidney;
Na_V_1.5: Voltage-gated cardiac sodium channel;
LQT: Long QT Syndrome;
VUS: Variant of Uncertain Significance

## Acknowledgements

We thank Kenneth Matreyek and Douglas Fowler for sharing HEK293 cell lines and Tim Strassmaier and Carlos Vanoye for helpful advice. The Nanion SyncroPatch 384PE is housed and managed within the Vanderbilt High-Throughput Screening center, an institutionally supported core, and was funded by NIH Shared Instrumentation Grant 1S10OD025281.

## Sources of Funding

This research was funded by F32 HL137385 (AMG), K99 HL135442 (BMK), and P50 GM115305 (DR).

## Disclosures

The authors have no conflicts to disclose.

## Literature Cited

1. Kapplinger JD, Tester DJ, Alders M, Benito B, Berthet M, Brugada J, Brugada P, Fressart V, Guerchicoff A, Harris-Kerr C, Kamakura S, Kyndt F, Koopmann TT, Miyamoto Y, Pfeiffer R, Pollevick GD, Probst V, Zumhagen S, Vatta M, Towbin JA, Shimizu W, Schulze-Bahr E, Antzelevitch C, Salisbury BA, Guicheney P, Wilde AAM, Brugada R, Schott JJ and Ackerman MJ. An international compendium of mutations in the SCN5A-encoded cardiac sodium channel in patients referred for Brugada syndrome genetic testing. Heart Rhythm. 2010;7:33–46.

2. Kapplinger JD, Tester DJ, Salisbury BA, Carr JL, Harris-Kerr C, Pollevick GD, Wilde AAM and Ackerman MJ. Spectrum and prevalence of mutations from the first 2,500 consecutive unrelated patients referred for the FAMILION (R) long QT syndrome genetic test. Heart Rhythm. 2009;6:1297–1303.

3. Moreau A, Gosselin-Badaroudine P, Delemotte L, Klein ML and Chahine M. Gating pore currents are defects in common with two Nav1.5 mutations in patients with mixed arrhythmias and dilated cardiomyopathy. J Gen Physiol. 2015;145:93–106.

4. Bezzina CR, Rook MB, Groenewegen WA, Herfst LJ, van der Wal AC, Lam J, Jongsma HJ, Wilde AA and Mannens MM. Compound heterozygosity for mutations (W156X and R225W) in SCN5A associated with severe cardiac conduction disturbances and degenerative changes in the conduction system. Circ Res. 2003;92:159–68.

5. Gui J, Wang T, Jones RP, Trump D, Zimmer T and Lei M. Multiple loss-of-function mechanisms contribute to SCN5A-related familial sick sinus syndrome. PLoS One. 2010;5:e10985.

6. Brugada J, Campuzano O, Arbelo E, Sarquella-Brugada G and Brugada R. Present Status of Brugada Syndrome: JACC State-of-the-Art Review. J Am Coll Cardiol. 2018;72:1046–1059.

7. Hosseini SM, Kim R, Udupa S, Costain G, Jobling R, Liston E, Jamal SM, Szybowska M, Morel CF, Bowdin S, Garcia J, Care M, Sturm AC, Novelli V, Ackerman MJ, Ware JS, Hershberger RE, Wilde AAM, Gollob MH and National Institutes of Health Clinical Genome Resource C. Reappraisal of Reported Genes for Sudden Arrhythmic Death. Circulation. 2018;138:1195–1205.

8. Milman A, Andorin A, Postema PG, Gourraud JB, Sacher F, Mabo P, Kim SH, Maeda S, Takahashi Y, Kamakura T, Aiba T, Conte G, Juang JJM, Leshem E, Michowitz Y, Fogelman R, Hochstadt A, Mizusawa Y, Giustetto C, Arbelo E, Huang Z, Corrado D, Delise P, Allocca G, Takagi M, Wijeyeratne YD, Mazzanti A, Brugada R, Casado-Arroyo R, Champagne J, Calo L, Sarquella-Brugada G, Jespersen CH, Tfelt-Hansen J, Veltmann C, Priori SG, Behr ER, Yan GX, Brugada J, Gaita F, Wilde AAM, Brugada P, Kusano KF, Hirao K, Nam GB, Probst V and Belhassen B. Ethnic differences in patients with Brugada syndrome and arrhythmic events: New insights from Survey on Arrhythmic Events in Brugada Syndrome. Heart Rhythm. 2019;16:1468–1474.

9. Kroncke BM, Glazer AM, Smith DK, Blume JD and Roden DM. SCN5A (NaV1.5) Variant Functional Perturbation and Clinical Presentation: Variants of a Certain Significance. Circ Genom Precis Med. 2018;11:e002095.

10. Wang DW, Yazawa K, George AL, Jr. and Bennett PB. Characterization of human cardiac Na+ channel mutations in the congenital long QT syndrome. Proc Natl Acad Sci U S A. 1996;93:13200–5.

11. Kalia SS, Adelman K, Bale SJ, Chung WK, Eng C, Evans JP, Herman GE, Hufnagel SB, Klein TE, Korf BR, McKelvey KD, Ormond KE, Richards CS, Vlangos CN, Watson M, Martin CL and Miller DT. Recommendations for reporting of secondary findings in clinical exome and genome sequencing, 2016 update (ACMG SF v2.0): a policy statement of the American College of Medical Genetics and Genomics. Genet Med. 2017;19:249–255.

12. Kroncke BM, Mendenhall J, Smith DK, Sanders CR, Capra JA, George AL, Blume JD, Meiler J and Roden DM. Protein structure aids predicting functional perturbation of missense variants in SCN5A and KCNQ1. Comput Struct Biotechnol J. 2019;17:206–214.

13. Kroncke BMS, D.; Glazer, A.; Roden, D.; Blume, J. A Bayesian method using sparse data to estimate penetrance of disease-associated genetic variants. bioRxiv: https://www.biorxivorg/content/101101/571158v1. 2019.

14. Richards S, Aziz N, Bale S, Bick D, Das S, Gastier-Foster J, Grody WW, Hegde M, Lyon E, Spector E, Voelkerding K, Rehm HL and Committee ALQA. Standards and guidelines for the interpretation of sequence variants: a joint consensus recommendation of the American College of Medical Genetics and Genomics and the Association for Molecular Pathology. Genet Med. 2015;17:405–24.

15. Kang SV, C.; Misra, S.; Echevarria, D.; Calhoun, J.; O’Connor, J.; Fabre, K.; McKnight, D.; Demmer, L.; Goldenberg, P.; Grote, L.; Thiffault, I.; Saunders, C.; Strauss, K.; Torkamani A.; van der Smagt, J.; van Gassen, K.; Cardon, R.; Diaz, J.; Leon, E.; Jacher, J.; Hannibal, M.; Litwin, J.; Friedman, N.; Schreiber, A.; Lynch, B.; Poduri, A.; Marsh, E.; Goldberg, E.; Millichap, J.; George, A.; Kearney, J. Spectrum of KV2.1 dysfunction in KCNB1-associated neurodevelopmental disorders. Annals of Neurology (in press). 2019.

16. Ng CP, M.; Liang, W.; Smith, N.; Foo, B.; Shrier, A.; Lukacs, G.; Hill, A.; Vandenberg, J. High-throughput phenotyping of heteromeric human ether-à-go-go-related gene potassium channel variants can discriminate pathogenic from rare benign variants. Heart Rhythm. 2019.

17. Vanoye CG, Desai RR, Fabre KL, Gallagher SL, Potet F, DeKeyser JM, Macaya D, Meiler J, Sanders CR and George AL, Jr. High-Throughput Functional Evaluation of KCNQ1 Decrypts Variants of Unknown Significance. Circ Genom Precis Med. 2018;11:e002345.

18. Lek M, Karczewski KJ, Minikel EV, Samocha KE, Banks E, Fennell T, O’Donnell-Luria AH, Ware JS, Hill AJ, Cummings BB, Tukiainen T, Birnbaum DP, Kosmicki JA, Duncan LE, Estrada K, Zhao F, Zou J, Pierce-Hoffman E, Berghout J, Cooper DN, Deflaux N, DePristo M, Do R, Flannick J, Fromer M, Gauthier L, Goldstein J, Gupta N, Howrigan D, Kiezun A, Kurki MI, Moonshine AL, Natarajan P, Orozco L, Peloso GM, Poplin R, Rivas MA, Ruano-Rubio V, Rose SA, Ruderfer DM, Shakir K, Stenson PD, Stevens C, Thomas BP, Tiao G, Tusie-Luna MT, Weisburd B, Won HH, Yu D, Altshuler DM, Ardissino D, Boehnke M, Danesh J, Donnelly S, Elosua R, Florez JC, Gabriel SB, Getz G, Glatt SJ, Hultman CM, Kathiresan S, Laakso M, McCarroll S, McCarthy MI, McGovern D, McPherson R, Neale BM, Palotie A, Purcell SM, Saleheen D, Scharf JM, Sklar P, Sullivan PF, Tuomilehto J, Tsuang MT, Watkins HC, Wilson JG, Daly MJ, MacArthur DG and Exome Aggregation C. Analysis of protein-coding genetic variation in 60,706 humans. Nature. 2016;536:285–91.

19. Matreyek KA, Stephany JJ and Fowler DM. A platform for functional assessment of large variant libraries in mammalian cells. Nucleic Acids Res. 2017;45:e102.

20. Matreyek KAS, J.; Chiasson, M.; Hasle, N.; Fowler, D. An Improved Platform for Functional Assessment of Large Protein Libraries in Mammalian Cells. Nucleic Acids Res. 2019.

21. Hermann M, Stillhard P, Wildner H, Seruggia D, Kapp V, Sanchez-Iranzo H, Mercader N, Montoliu L, Zeilhofer HU and Pelczar P. Binary recombinase systems for high-resolution conditional mutagenesis. Nucleic Acids Res. 2014;42:3894–907.

22. Clatot J, Ziyadeh-Isleem A, Maugenre S, Denjoy I, Liu H, Dilanian G, Hatem SN, Deschenes I, Coulombe A, Guicheney P and Neyroud N. Dominant-negative effect of SCN5A N-terminal mutations through the interaction of Na(v)1.5 alpha-subunits. Cardiovasc Res. 2012;96:53–63.

23. Makiyama T, Akao M, Tsuji K, Doi T, Ohno S, Takenaka K, Kobori A, Ninomiya T, Yoshida H, Takano M, Makita N, Yanagisawa F, Higashi Y, Takeyama Y, Kita T and Horie M. High risk for bradyarrhythmic complications in patients with Brugada syndrome caused by SCN5A gene mutations. J Am Coll Cardiol. 2005;46:2100–6.

24. Pfahnl AE, Viswanathan PC, Weiss R, Shang LL, Sanyal S, Shusterman V, Kornblit C, London B and Dudley SC, Jr. A sodium channel pore mutation causing Brugada syndrome. Heart Rhythm. 2007;4:46–53.

25. Valdivia CR, Ackerman MJ, Tester DJ, Wada T, McCormack J, Ye B and Makielski JC. A novel SCN5A arrhythmia mutation, M1766L, with expression defect rescued by mexiletine. Cardiovasc Res. 2002;55:279–89.

26. Valdivia CR, Tester DJ, Rok BA, Porter CB, Munger TM, Jahangir A, Makielski JC and Ackerman MJ. A trafficking defective, Brugada syndrome-causing SCN5A mutation rescued by drugs. Cardiovasc Res. 2004;62:53–62.

27. Whiffin N, Minikel E, Walsh R, O’Donnell-Luria AH, Karczewski K, Ing AY, Barton PJR, Funke B, Cook SA, MacArthur D and Ware JS. Using high-resolution variant frequencies to empower clinical genome interpretation. Genet Med. 2017;19:1151–1158.

28. Choi Y and Chan AP. PROVEAN web server: a tool to predict the functional effect of amino acid substitutions and indels. Bioinformatics. 2015;31:2745–7.

29. Adzhubei IA, Schmidt S, Peshkin L, Ramensky VE, Gerasimova A, Bork P, Kondrashov AS and Sunyaev SR. A method and server for predicting damaging missense mutations. Nat Methods. 2010;7:248–249.

30. Landrum MJ, Lee JM, Benson M, Brown G, Chao C, Chitipiralla S, Gu B, Hart J, Hoffman D, Hoover J, Jang W, Katz K, Ovetsky M, Riley G, Sethi A, Tully R, Villamarin-Salomon R, Rubinstein W and Maglott DR. ClinVar: public archive of interpretations of clinically relevant variants. Nucleic Acids Res. 2016;44:D862–8.

31. Leaver-Fay A, Tyka M, Lewis SM, Lange OF, Thompson J, Jacak R, Kaufman K, Renfrew PD, Smith CA, Sheffler W, Davis IW, Cooper S, Treuille A, Mandell DJ, Richter F, Ban YE, Fleishman SJ, Corn JE, Kim DE, Lyskov S, Berrondo M, Mentzer S, Popovic Z, Havranek JJ, Karanicolas J, Das R, Meiler J, Kortemme T, Gray JJ, Kuhlman B, Baker D and Bradley P. ROSETTA3: an object-oriented software suite for the simulation and design of macromolecules. Methods Enzymol. 2011;487:545–74.

32. Shen HZ, Liu DL, Wu K, Lei JL and Yan N. Structures of human Na(v)1.7 channel in complex with auxiliary subunits and animal toxins. Science. 2019;363:1303–1308.

33. Shen H, Li Z, Jiang Y, Pan X, Wu J, Cristofori-Armstrong B, Smith JJ, Chin YKY, Lei J, Zhou Q, King GF and Yan N. Structural basis for the modulation of voltage-gated sodium channels by animal toxins. Science. 2018;362.

34. Itoh H, Berthet M, Fressart V, Denjoy I, Maugenre S, Klug D, Mizusawa Y, Makiyama T, Hofman N, Stallmeyer B, Zumhagen S, Shimizu W, Wilde AA, Schulze-Bahr E, Horie M, Tezenas du Montcel S and Guicheney P. Asymmetry of parental origin in long QT syndrome: preferential maternal transmission of KCNQ1 variants linked to channel dysfunction. Eur J Hum Genet. 2016;24:1160–6.

35. Yue P, Li Z and Moult J. Loss of protein structure stability as a major causative factor in monogenic disease. J Mol Biol. 2005;353:459–73.

36. Stein A, Fowler DM, Hartmann-Petersen R and Lindorff-Larsen K. Biophysical and Mechanistic Models for Disease-Causing Protein Variants. Trends Biochem Sci. 2019;44:575–588.

37. Park H, Bradley P, Greisen P, Liu Y, Mulligan VK, Kim DE, Baker D and DiMaio F. Simultaneous Optimization of Biomolecular Energy Functions on Features from Small Molecules and Macromolecules. Journal of Chemical Theory and Computation. 2016;12:6201–6212.

38. Kim MK and Kang YK. Positional preference of proline in alpha-helices. Protein Sci. 1999;8:1492–9.

39. Milman A, Gourraud JB, Andorin A, Postema PG, Sacher F, Mabo P, Conte G, Giustetto C, Sarquella-Brugada G, Hochstadt A, Kim SH, Juang JJM, Maeda S, Takahashi Y, Kamakura T, Aiba T, Leshem E, Michowitz Y, Rahkovich M, Mizusawa Y, Arbelo E, Huang Z, Denjoy I, Wijeyeratne YD, Napolitano C, Brugada R, Casado-Arroyo R, Champagne J, Calo L, Tfelt-Hansen J, Priori SG, Takagi M, Veltmann C, Delise P, Corrado D, Behr ER, Gaita F, Yan GX, Brugada J, Leenhardt A, Wilde AAM, Brugada P, Kusano KF, Hirao K, Nam GB, Probst V and Belhassen B. Gender differences in patients with Brugada syndrome and arrhythmic events: Data from a survey on arrhythmic events in 678 patients. Heart Rhythm. 2018;15:1457–1465.

40. Bezzina CR, Barc J, Mizusawa Y, Remme CA, Gourraud JB, Simonet F, Verkerk AO, Schwartz PJ, Crotti L, Dagradi F, Guicheney P, Fressart V, Leenhardt A, Antzelevitch C, Bartkowiak S, Schulze-Bahr E, Zumhagen S, Behr ER, Bastiaenen R, Tfelt-Hansen J, Olesen MS, Kaab S, Beckmann BM, Weeke P, Watanabe H, Endo N, Minamino T, Horie M, Ohno S, Hasegawa K, Makita N, Nogami A, Shimizu W, Aiba T, Froguel P, Balkau B, Lantieri O, Torchio M, Wiese C, Weber D, Wolswinkel R, Coronel R, Boukens BJ, Bezieau S, Charpentier E, Chatel S, Despres A, Gros F, Kyndt F, Lecointe S, Lindenbaum P, Portero V, Violleau J, Gessler M, Tan HL, Roden DM, Christoffels VM, Le Marec H, Wilde AA, Probst V, Schott JJ, Dina C, Redon R, Borggrefe M and Schimpf R. Common variants at SCN5A-SCN10A and HEY2 are associated with Brugada syndrome, a rare disease with high risk of sudden cardiac death (vol 45, pg 1044, 2013). Nat Genet. 2013;45:1409–1409.

41. Takehara N, Makita N, Kawabe J, Sato N, Kawamura Y, Kitabatake A and Kikuchi K. A cardiac sodium channel mutation identified in Brugada syndrome associated with atrial standstill. J Intern Med. 2004;255:137–42.

42. Probst V, Allouis M, Sacher F, Pattier S, Babuty D, Mabo P, Mansourati J, Victor J, Nguyen JM, Schott JJ, Boisseau P, Escande D and Le Marec H. Progressive cardiac conduction defect is the prevailing phenotype in carriers of a Brugada syndrome SCN5A mutation. J Cardiovasc Electrophysiol. 2006;17:270–5.

43. Remme CA, Wilde AA and Bezzina CR. Cardiac sodium channel overlap syndromes: different faces of SCN5A mutations. Trends Cardiovasc Med. 2008;18:78–87.

44. Frigo G, Rampazzo A, Bauce B, Pilichou K, Beffagna G, Danieli GA, Nava A and Martini B. Homozygous SCN5A mutation in Brugada syndrome with monomorphic ventricular tachycardia and structural heart abnormalities. Europace. 2007;9:391–7.

45. Benson DW, Wang DW, Dyment M, Knilans TK, Fish FA, Strieper MJ, Rhodes TH and George AL, Jr. Congenital sick sinus syndrome caused by recessive mutations in the cardiac sodium channel gene (SCN5A). J Clin Invest. 2003;112:1019–28.

46. Anderson CL, Delisle BP, Anson BD, Kilby JA, Will ML, Tester DJ, Gong Q, Zhou Z, Ackerman MJ and January CT. Most LQT2 mutations reduce Kv11.1 (hERG) current by a class 2 (trafficking-deficient) mechanism. Circulation. 2006;113:365–73.

47. Casadio R, Vassura M, Tiwari S, Fariselli P and Luigi Martelli P. Correlating disease-related mutations to their effect on protein stability: a large-scale analysis of the human proteome. Hum Mutat. 2011;32:1161–70.

48. Zhou Y and Bowie JU. Building a thermostable membrane protein. J Biol Chem. 2000;275:6975–9.

49. Chavali NV, Kryshtal DO, Parikh SS, Wang L, Glazer AM, Blackwell DJ, Kroncke BM, Shoemaker MB and Knollmann BC. Patient-independent human induced pluripotent stem cell model: A new tool for rapid determination of genetic variant pathogenicity in long QT syndrome. Heart Rhythm. 2019.

50. Fatima A, Kaifeng S, Dittmann S, Xu G, Gupta MK, Linke M, Zechner U, Nguemo F, Milting H, Farr M, Hescheler J and Saric T. The disease-specific phenotype in cardiomyocytes derived from induced pluripotent stem cells of two long QT syndrome type 3 patients. PLoS One. 2013;8:e83005.

51. Selga E, Sendfeld F, Martinez-Moreno R, Medine CN, Tura-Ceide O, Wilmut SI, Perez GJ, Scornik FS, Brugada R and Mills NL. Sodium channel current loss of function in induced pluripotent stem cell-derived cardiomyocytes from a Brugada syndrome patient. J Mol Cell Cardiol. 2018;114:10–19.

52. Heyne HB-N, D.; Iqbal, S.; Palmer, D.; Brunklaus, A.; the Epi25 Collaborative; Johannesen, K.; Lauxmann, S.; Lemke, J.; Moller, R.; Perez-Palma, E.; Scholl, U.; Syrbe, S.; Lerche, H; May, P.; Lal, D.; Campbell, A.; Pan, J.; Wang, H.; Daly, M. Predicting functional effects of missense variants in the voltage-gated sodium and calcium channels. bioRxiv http://dxdoiorg/101101/671453. 2019.

53. Li B, Mendenhall JL, Kroncke BM, Taylor KC, Huang H, Smith DK, Vanoye CG, Blume JD, George AL, Jr., Sanders CR and Meiler J. Predicting the Functional Impact of KCNQ1 Variants of Unknown Significance. Circ Cardiovasc Genet. 2017;10.

54. Findlay GM, Daza RM, Martin B, Zhang MD, Leith AP, Gasperini M, Janizek JD, Huang X, Starita LM and Shendure J. Accurate classification of BRCA1 variants with saturation genome editing. Nature. 2018;562:217–222.

55. Glazer AK, B.; Matreyek, K.; Yang, T.; Wada, Y.; Shields, T.; Salem, J.; Fowler, D.; Roden, D. Deep Mutational Scan of a cardiac sodium channel voltage sensor. bioRxiv https://www.biorxivorg/content/101101/773887v1. 2019.

56. Matreyek KA, Starita LM, Stephany JJ, Martin B, Chiasson MA, Gray VE, Kircher M, Khechaduri A, Dines JN, Hause RJ, Bhatia S, Evans WE, Relling MV, Yang WJ, Shendure J and Fowler DM. Multiplex assessment of protein variant abundance by massively parallel sequencing. Nat Genet. 2018;50:874-+.

57. Aiba T, Farinelli F, Kostecki G, Hesketh GG, Edwards D, Biswas S, Tung L and Tomaselli GF. A mutation causing Brugada syndrome identifies a mechanism for altered autonomic and oxidant regulation of cardiac sodium currents. Circ Cardiovasc Genet. 2014;7:249–56.

58. Casini SA, M.; Wang, Z.; Portero, V.; Ross-Kaschitza, D.; Rougier, J.; Marchal, G.; Chung, W.; Bezzina, C.; Abriel, H.; Remme, C. Functional Consequences of the SCN5A-p.Y1977N Mutation within the PY Ubiquitylation Motif: Discrepancy between HEK293 Cells and Transgenic Mice. Int J Mol Sci. 2019;11.

59. Clatot J, Hoshi M, Wan X, Liu H, Jain A, Shinlapawittayatorn K, Marionneau C, Ficker E, Ha T and Deschenes I. Voltage-gated sodium channels assemble and gate as dimers. Nat Commun. 2017;8:2077.

60. Makielski JC, Ye B, Valdivia CR, Pagel MD, Pu J, Tester DJ and Ackerman MJ. A ubiquitous splice variant and a common polymorphism affect heterologous expression of recombinant human SCN5A heart sodium channels. Circ Res. 2003;93:821–8.

