## Supplement for "High-throughput reclassification of *SCN5A* variants"

### Table of Contents

**Supplemental Methods**

**Figure S1. Diagram of cloning and cell line generation**
**Figure S2. Voltage protocols used in this study**
**Figure S3. Example sodium current traces from included and excluded wells**
**Figure S4: Classification criteria**
**Figure S5. A structural model of human SCN5A bound with SCN1B.**
**Figure S6. Voltage dependence of activation**
**Figure S7. Inactivation time**
**Figure S8. Voltage Dependence of Inactivation**
**Figure S9. Recovery from inactivation**
**Figure S10. Late current**
**Figure S11. Variant distance from the pore is strongly correlated with normalized peak current density**
**Figure S12. Variant-induced change in thermostability is correlated with functional impact.**
**Figure S13. Variants may compromise function by disrupting the pore**

**Table S1. Patient and gnomAD counts, peak current density, and classifications**
**Table S2. Zone boundaries and restriction enzymes**
**Table S3. Primers used in this study**
**Table S4: Summary of the Rosetta energy functions used for $\Delta\Delta G$ calculations**
**Table S5. All measured parameters**
**Table S6. *SCN5A* missense variants with <10% peak current density**

**File S1. Summary of patch clamp data for each variant (.csv)**

**Supplemental Methods**

Peak current averaging

For each variant, mean peak current density was calculated as *peak current/capacitance*, averaged across all in voltage control cells. For variants with near-wildtype peak currents, the percentage of cells with detectable currents matched the expected percentage from flow cytometry (Figure 1A-B). However, severe loss of function variants had a substantially lower proportion of wells with detectable currents compared to the predicted percentage from flow cytometry, as is typically observed.[1] Since a simple mean would overestimate the true peak current, an adjustment was made to the means based on a comparison between the flow cytometry and patch clamp data. If the percentage of current-positive wells was >10% less than the expected percentage from flow cytometry, the number of expected additional cells with no current was calculated as follows:

*Expected additional cells = #QC+ cells *(%mCherry+ cells - %QC+ cells with current)*

QC+ cells are defined as cells with a 0.5- 10 GΩ seal resistance and 5-30 pF capacitance. These expected additional cells were added to the peak current density average. For each variant, peak current density averages were first normalized to wildtype cells from the same experiment/transfection, then averaged across independent experiments/transfections.

Building a homology model

Two starting partial models of SCN5A bound with SCN1B, which only covered aligned residues, were generated by threading the sequences of SCN5A and SCN1B onto the two template structures respectively. The threading was guided by the corresponding sequence alignments. Full models were created by hybridization of the two starting models using the

Rosetta comparative modeling (RosettaCM) protocol[2] guided by the RosettaMembrane energy function.[3] The starting model generated from the primary template was used as the base model during hybridization. The boundaries of membrane-spanning segments were calculated using the PPM server[4] based on the starting model generated from the primary template. The boundaries were used to impose membrane-specific Rosetta energy terms on residues within the theoretical membrane bilayer. Fragments in the starting model where coordinates were missing were modeled *de novo* by inserting fragments selected from the Protein Data Bank using local sequence information.[5] Amino acid rotamer conformations were optimized by a nondeterministic Monte Carlo simulated annealing protocol, referred to as rotamer repacking in Rosetta, and models were refined in internal and Cartesian coordinate space by gradient-based minimization. A total of 6000 full models of SCN5A bound with SCN1B were generated using RosettaCM and models ranked by Rosetta energy function in the top 20% were grouped into ten clusters based on pairwise root-mean-square distances. The lowest-energy model from the largest cluster was selected as the final model (Figure 5, S5) for structure-based analysis in this work. This model was also the lowest-energy model across all models sampled.

<u>ΔΔG Calculations</u>

The Rosetta ΔΔG protocol samples conformational degrees of freedom in a locally restricted region around the residue of interest: all sidechains within 6 Å of the mutated residue, and the backbone of a three-residue window around the mutated residue were allowed to move.[6] The sampling was guided by an energy function fitted to recapitulate experimentally determined membrane protein ΔΔG values (Table S4).[7] ΔΔG values computed through this energy function were previously demonstrated to be strongly correlated with cell surface expression levels of human rhodopsin variants.[8] Previous studies have shown that the magnitude of variant-caused perturbation to protein native thermostability (i.e. $|\Delta\Delta G|$) correlates best with disease likelihood.[9] Accordingly, in this work, variant-induced perturbation to native thermostability ($|\Delta\Delta G_{wildtype\text{->}variant}|$) was computed as the absolute energy difference between the refined variant structure and the refined wild-type structure: $|\Delta\Delta G_{wildtype\text{->}variant}| = |\Delta G_{variant} - \Delta G_{wildtype}|$. Due to the nondeterministic nature of conformation sampling in Rosetta, the $|\Delta\Delta G|$ of each variant was computed 30 times and the average was recorded. The Rosetta energy function is a hybrid of both physically meaningful terms and statistics-based terms, and it does not compute the entropic contribution to the Gibbs free energy in a thermodynamically rigorous manner.[10] Thus, it is best to interpret $\Delta\Delta G$ values computed through the Rosetta energy function in a statistical manner as approximations to the Gibbs free energies.

**Figure S1. Diagram of cloning and cell line generation**

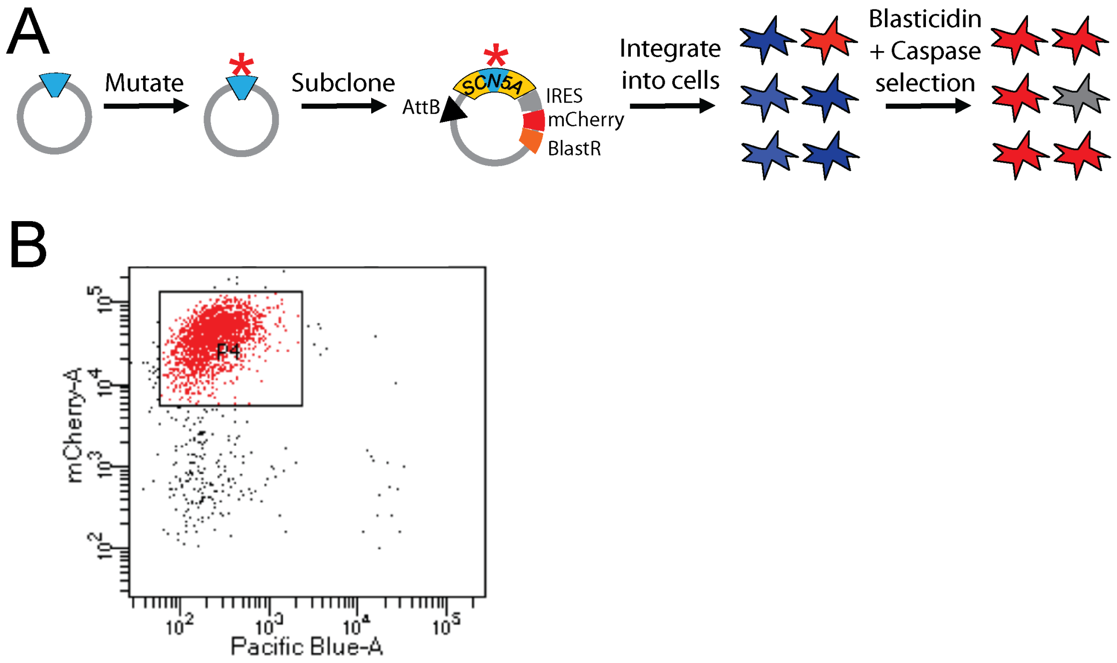

**Figure S2. Voltage protocols used in this study**

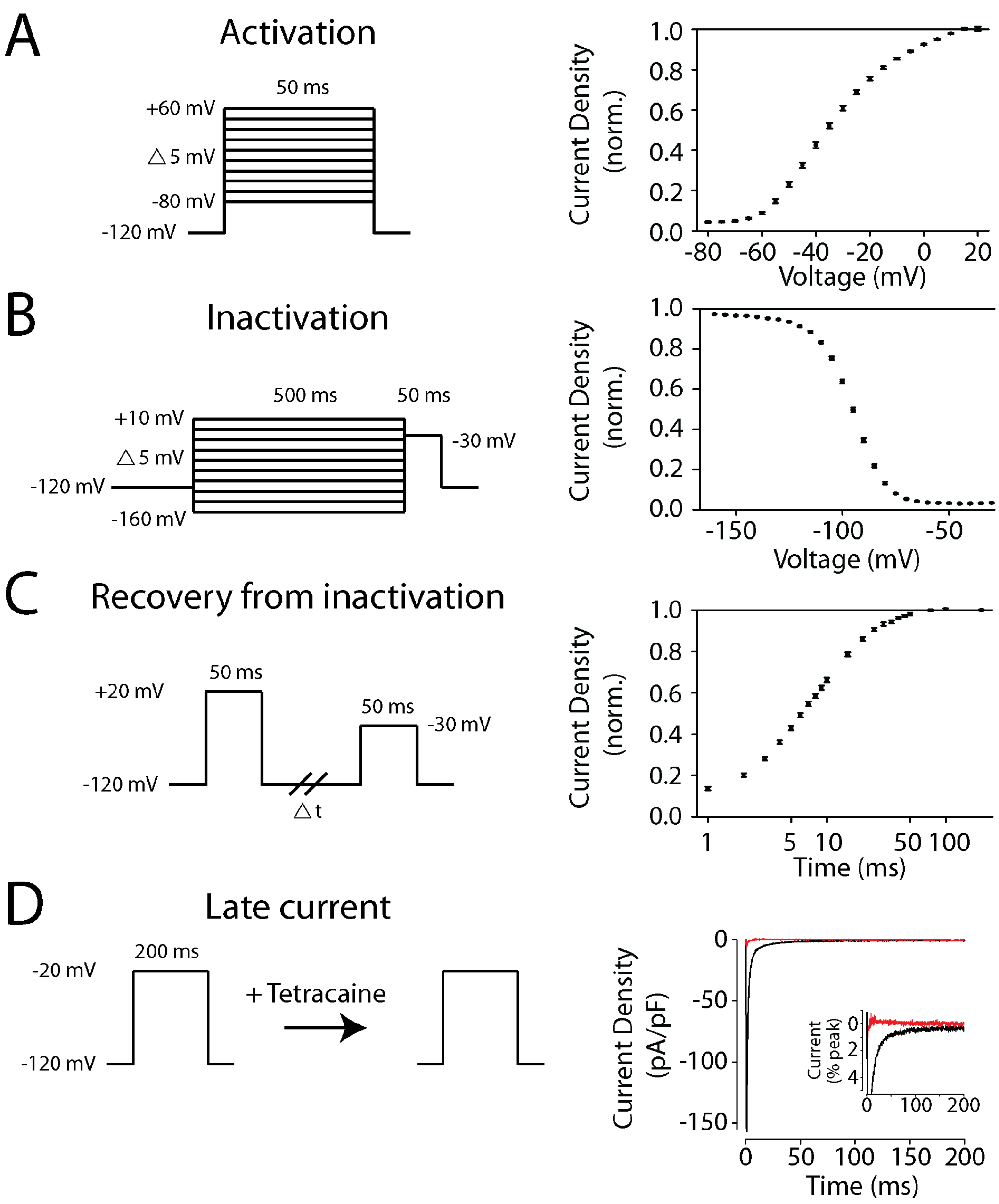

**Figure S3. Example sodium current traces from included and excluded wells**

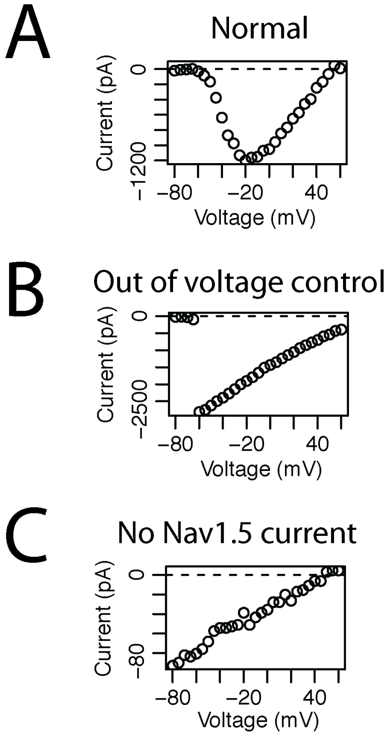

A) Example well with $Na_V1.5$ current that would be included in the analysis. B) Example well with $Na_V1.5$ current that is out of voltage control. This well would be excluded from the analysis. C) Example well with no substantial $Na_V1.5$ current. Note the difference in y-axis scale between panels.

**Figure S4: Classification criteria**

| | Benign: Strong | Benign: Supporting | Pathogenic: Supporting | Pathogenic: Moderate | Pathogenic: Strong | Pathogenic: Very Strong |
|---|---|---|---|---|---|---|
| **Population data** | MAF too high for disorder BA1/BS1 | | | Absent in population databases PM2 | Prevalence in cases statistically increased over controls PS4 | |
| **Computational data** | | Multiple lines of comp. evidence suggest no impact BP4 | Multiple lines of comp. evidence support deleterious effect PP3 | Novel missense at aa with diff. pathogenic missense PM5 | Same aa change as established pathogenic variant PS1 | Predicted null if LOF is known disease mechanism PVS1 |
| **Functional data** | Well-established functional studies show no deleterious effect BS3 | | Missense in gene with low rate of benign missense PP2 | Mutational hot spot or well-studied functional domain PM1 | Well-established functional studies show a deleterious effect PS3 | |
| **Segregation data** | Nonsegregation with disease BS4 | | Cosegregation with disease in mult. affected family members PP1 | Increased segregation data → | → | |
| **Other database** | | Reputable source reports benign BP6 | Reputable source reports pathogenic PP5 | | | |

Diagram of American College of Medical Genetics and Genomics classification criteria, with a focus on criteria relevant for classification of *SCN5A* missense variants. Figure is adapted from Richards et al, 2015.[11]

**Figure S5. A structural model of human SCN5A bound with SCN1B.**

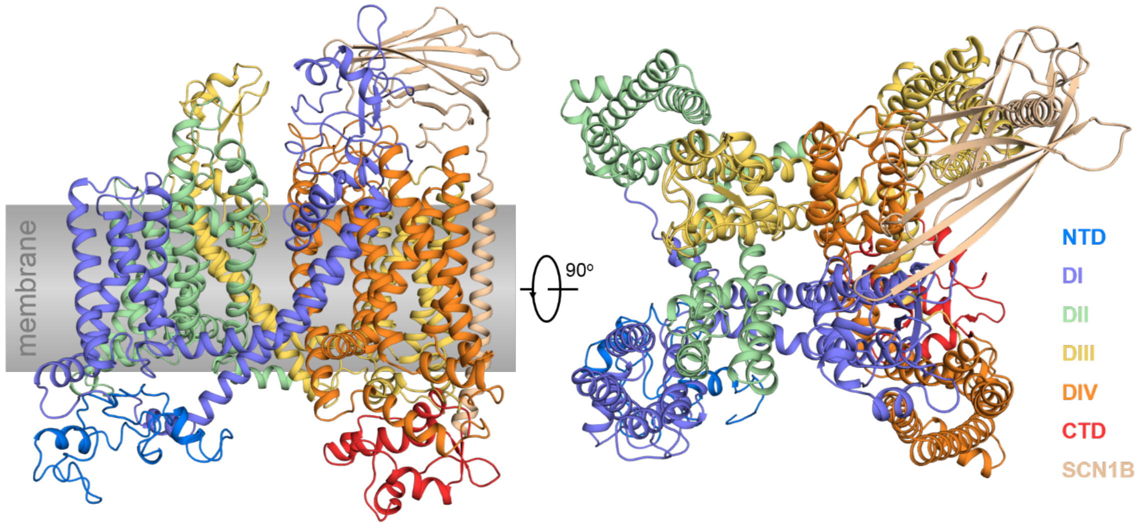

The N-terminal domain (NTD), four homologous ion channel domains (DI through DIV) each consisting of six transmembrane helices, C-terminal domain (CTD), and SCN1B are color-coded.

**Figure S6: Voltage Dependence of Activation**

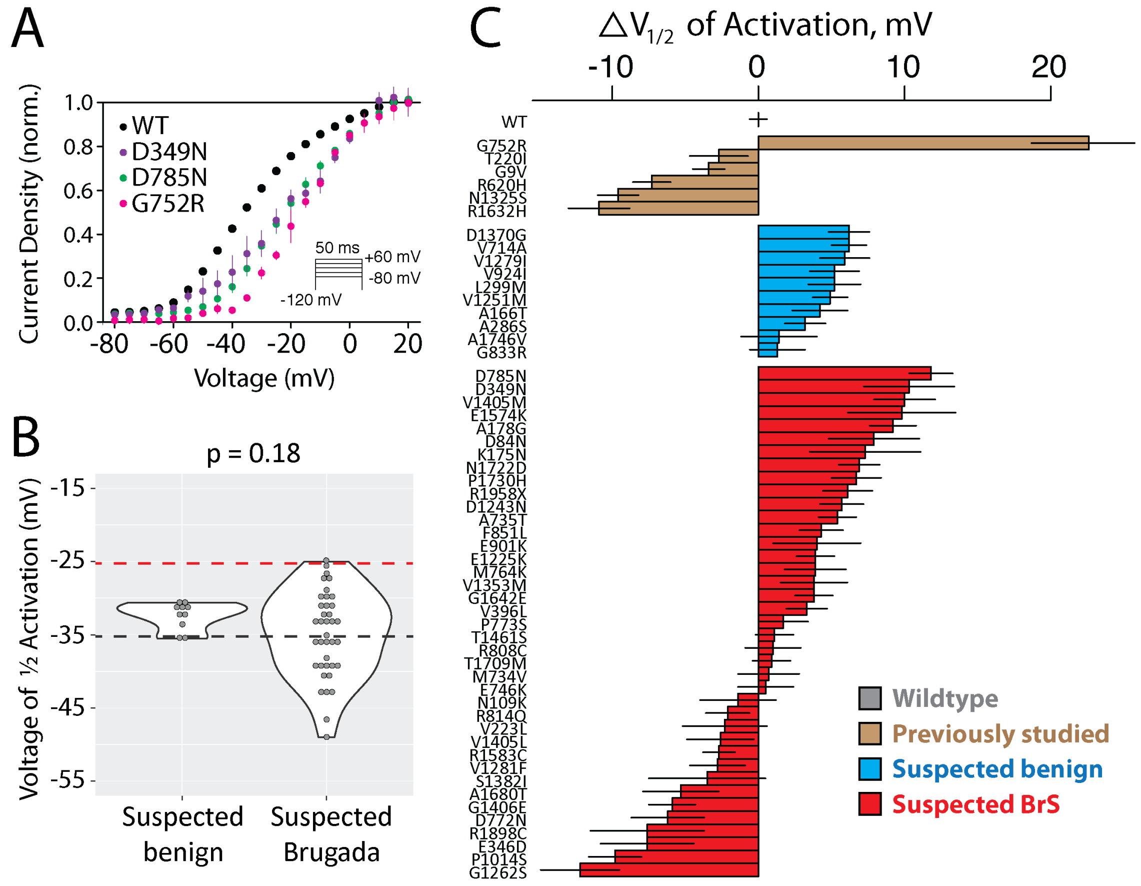

A) Normalized activation curves for wildtype, D785N and D349N (the two suspected Brugada Syndrome-associated variants with >10 mV rightward shifts), and G752R, a previously studied variant with a large rightward shift in $V_{1/2}$ activation, as has previously been observed.[12] All three of these variants also have a reduction in peak current (Figure 2). B) Violin plot of $V_{1/2}$ activation. Black line indicates wildtype value and red line indicates a 10 mV shift in activation. Two variants, D785N and D349N, have >10 mV rightward shift in activation voltage. Panel same as Figure 2E. C) Barplot of wildtype (grey), previously studied (brown), suspected benign (blue), or suspected Brugada Syndrome-associated voltage of ½ activation. Bars indicate mean

+/- standard errors. Only variants with at least 5 qualifying cells were included, so most severe loss of function variants are not included.

**Figure S7: Inactivation time**

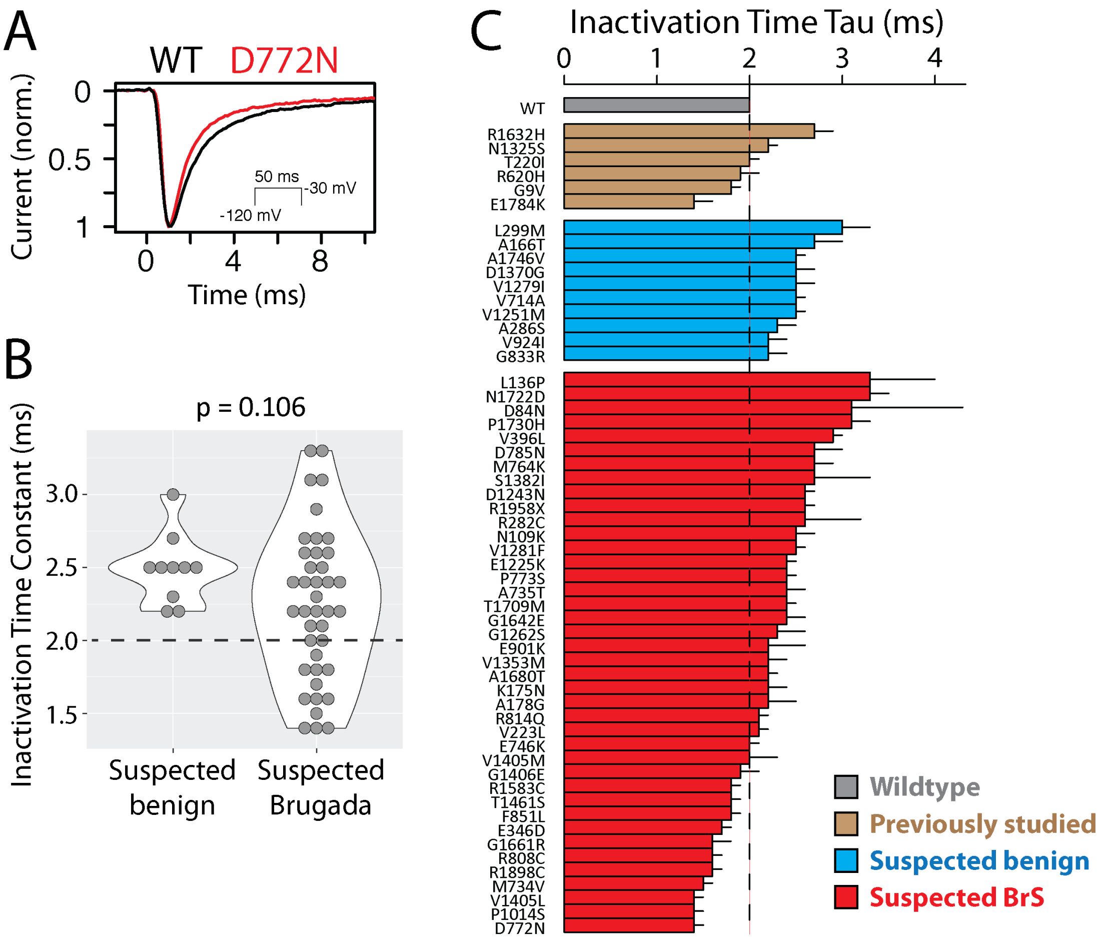

A) Inactivation time for wildtype and the suspected Brugada Syndrome-associated variant with the shortest time constant of inactivation, D772N. Inactivation time was measured from a 50ms pulse from -120 mV to -30 mV. Representative traces were chosen that had inactivation time constants closest to the mean values. B) Violin plot of inactivation time constant. Black line indicates wildtype value. C) Barplot of wildtype (grey), previously studied (brown), suspected benign (blue), or suspected Brugada Syndrome-associated inactivation time. Bars indicate mean +/- standard errors. Only variants with at least 5 qualifying cells were included, so most severe loss of function variants are not included.

**Figure S8: Voltage Dependence of Inactivation**

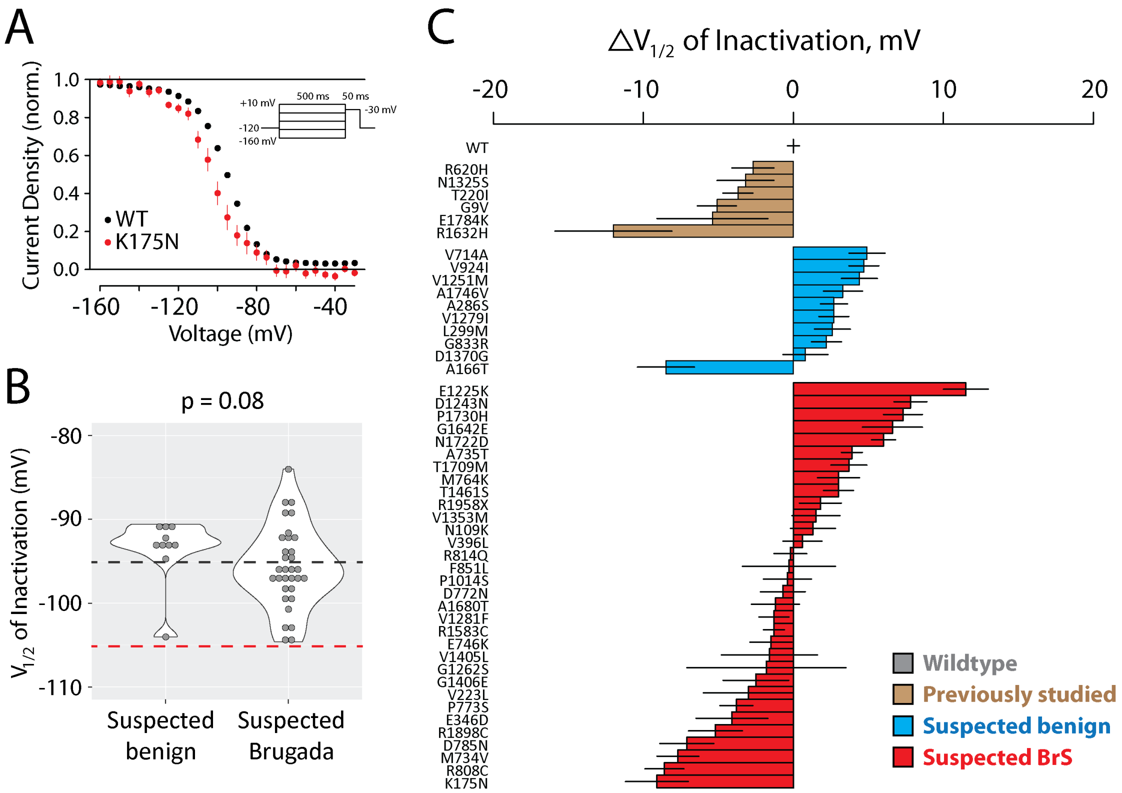

A) Normalized inactivation curves for wildtype and K175N, a suspected Brugada syndrome-associated variant that has the largest observed leftward shift in $V_{1/2}$ inactivation. B) Violin plot of $V_{1/2}$ inactivation. Black line indicates wildtype value and red line indicates a 10 mV shift in inactivation. No previously unstudied variants had a >10 mV leftward shift in inactivation voltage. C) Barplot of wildtype (grey), previously studied (brown), suspected benign (blue), or suspected Brugada Syndrome-associated voltage of ½ inactivation. Bars indicate mean +/- standard errors. Only variants with at least 5 qualifying cells were included, so most severe loss of function variants are not included.

**Figure S9: Recovery from Inactivation**

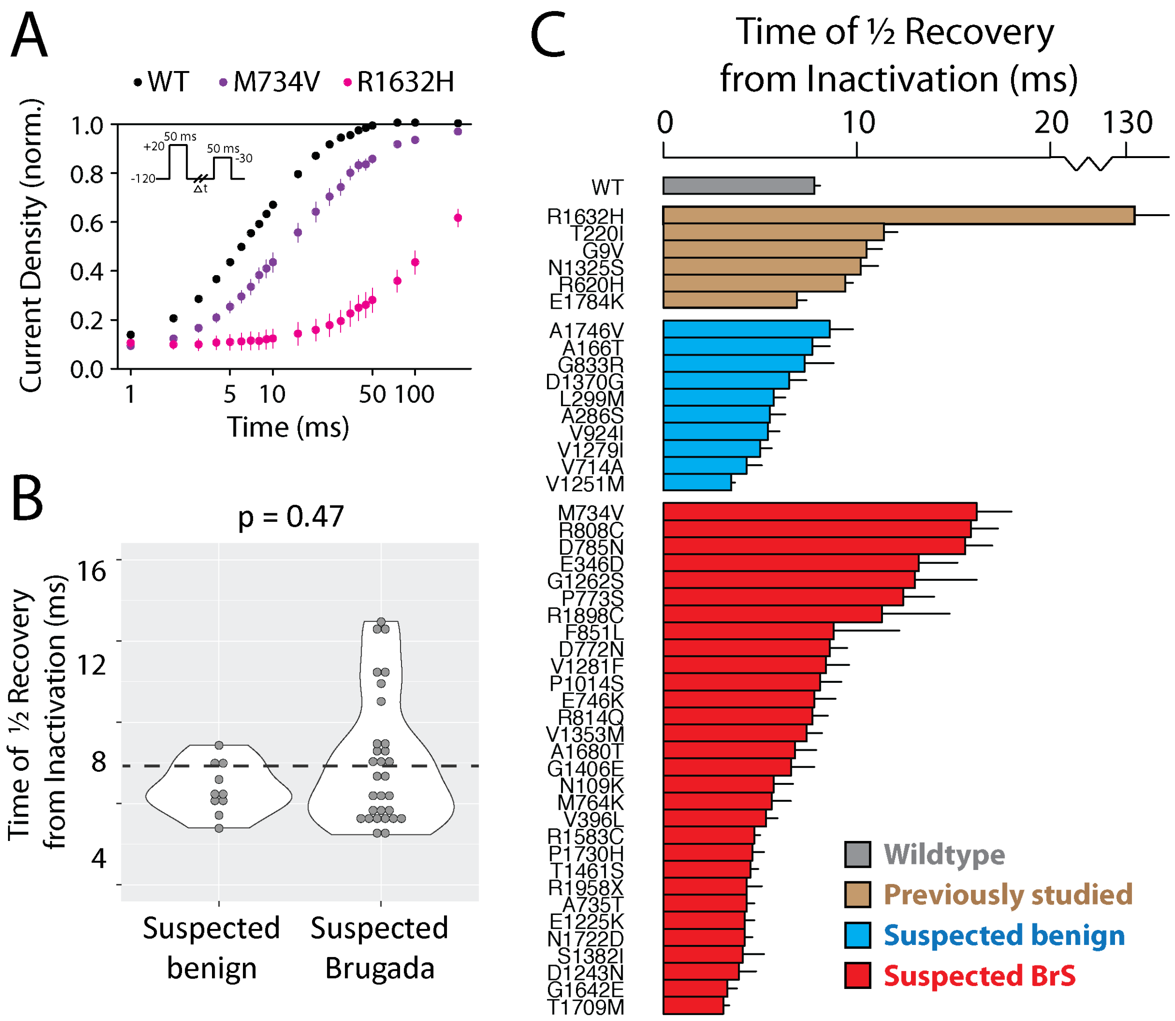

A) Normalized recovery inactivation curves for wildtype, M734V, the suspected Brugada syndrome-associated variant with the largest increase in recovery from inactivation time, and R1632H, a previously studied variant that has a very large recovery from inactivation.[13] The X-axis has a break to accommodate the large value for R1632H. B) Violin plot of time of ½ recovery from inactivation, derived from fitting an exponential fit to the data derived from each cell. Black line indicates wildtype value. C) Barplot of wildtype (grey), previously studied (brown), suspected benign (blue), or suspected Brugada Syndrome-associated time of ½ recovery from inactivation. Bars indicate mean +/- standard errors. Only variants with at least 5 qualifying cells were included, so most severe loss of function variants are not included.

**Figure S10: Late current**

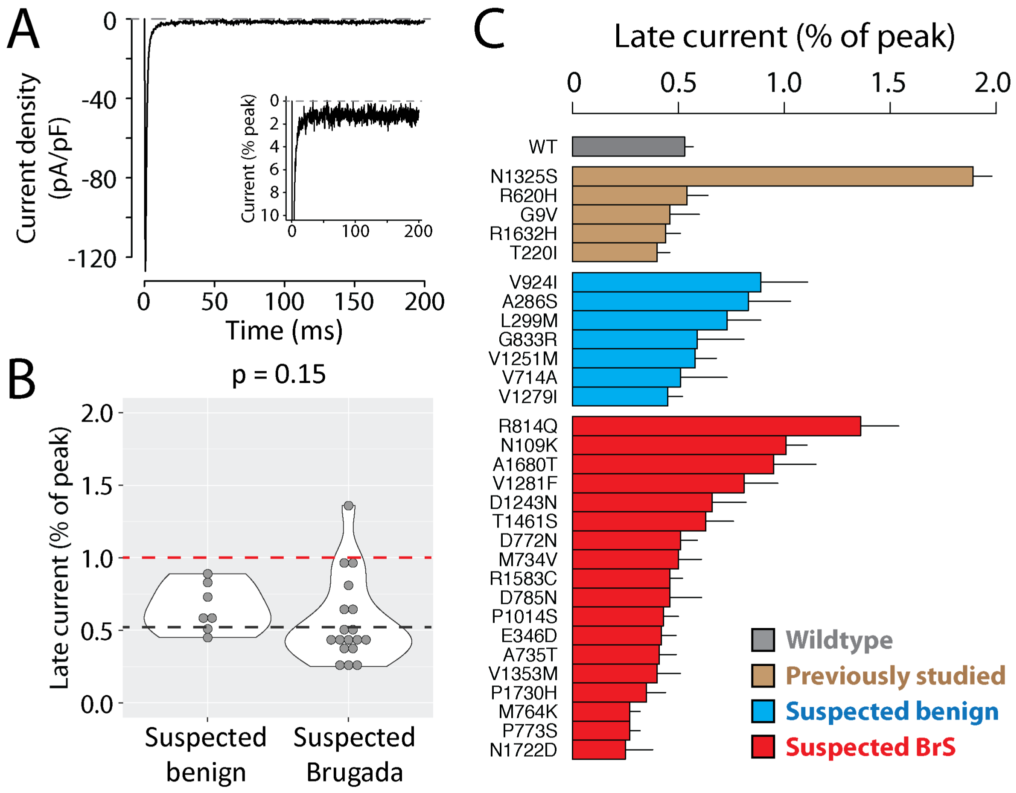

A) Tetracaine-sensitive late currents for R814Q, the suspected Brugada syndrome-associated variant that has the largest observed increase in late current. B) Violin plot of time of late current (normalized as the percentage of peak current). Black line indicates wildtype value and red line indicates the cutoff (1%) that was considered to be deleterious. C) Barplot of wildtype (grey), previously studied (brown), suspected benign (blue), or suspected Brugada Syndrome-associated late current (normalized as the percentage of peak current). Bars indicate mean +/- standard errors. Only variants with at least 5 qualifying cells were included, so most severe loss of function variants are not included.

**Figure S11: Variant distance from the pore is strongly correlated with normalized peak current density**

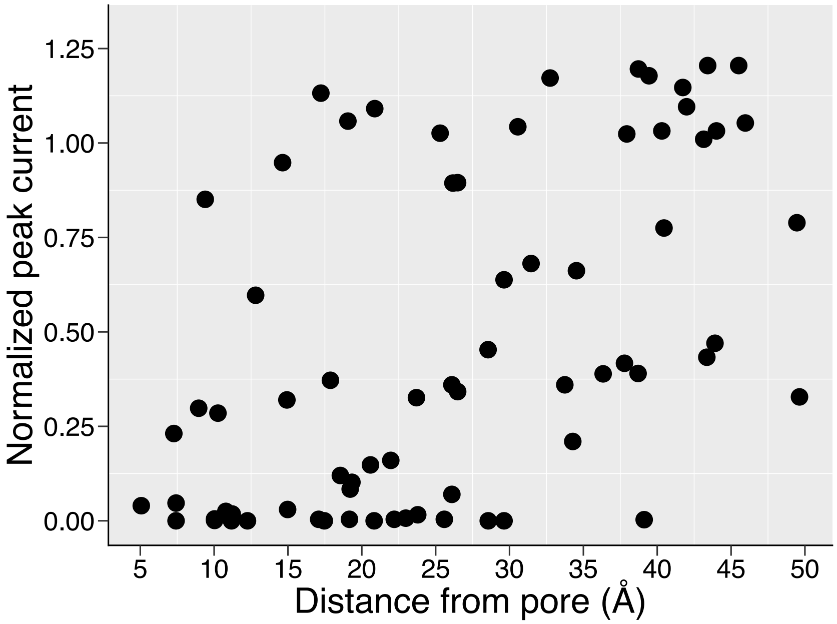

A scatter plot showing the correlation between variant distance from the pore and normalized peak currents (Pearson's $r = 0.538$, $p = 1.33\text{e} - 6$).

**Figure S12. Variant-induced perturbation to native thermostability is correlated with functional impact.**

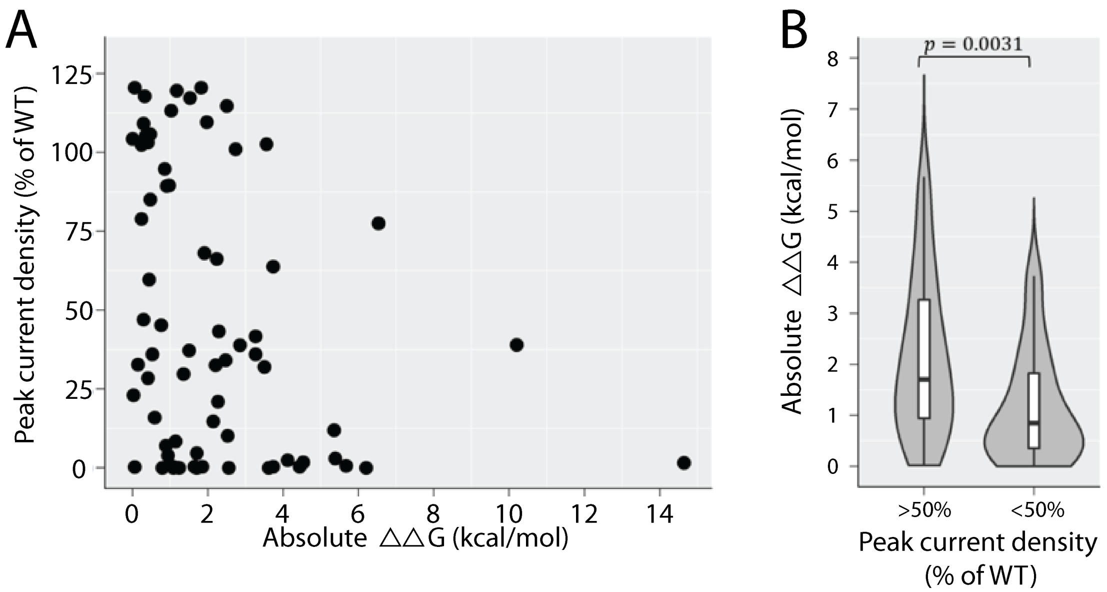

(A) A scatter plot showing the correlation between variant-induced perturbation to native thermostability (|ΔΔG|) and normalized peak currents (Pearson's $r = -0.31, p = 0.0092$). (B) A violin plot illustrating the distribution of absolute ΔΔG values of variants affecting SCN5A function (normalized peak current <50%, $n = 44$, median $|\Delta\Delta G| = 2.00\ kcal/mol$ ) and normal variants (normalized peak current >=50%, n=26 median $|\Delta\Delta G| = 0.88\ kcal/mol$, Mann-Whitney U test, $p = 0.0031$).

**Figure S13. Variants may compromise function by disrupting the pore**

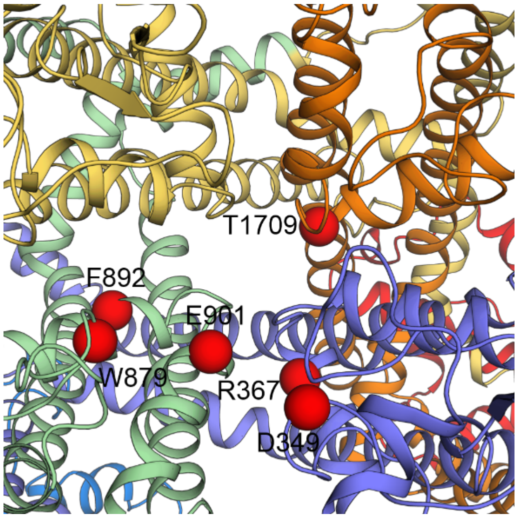

Seven pore-lining variants evaluated in this work, namely D349N, R367C, R367L, W879R, F892I, E901K, and T1709M, cause (partial) loss of function (<50% peak current), potentially by disrupting the pore. These variants induce only minor changes in perturbations to native thermostability. The absolute ΔΔG values of these seven variants are 0.41, 1.06, 1.21, 1.09, 0.79, 0.94, 0.03 kcal/mol and the normalized peak currents are 0.29, 0.01, 0.00, 0.00, 0.00, 0.04, and 0.23, respectively. Domains are color coded as in Fig. S4. The Cα atoms of mutated residues are rendered as red spheres. E901K is further highlighted in Figure 5D.

### Table S1. Patient and gnomAD counts, peak current density, and classifications

| Class | Variant | BrS1 | LQT3 | Unaff. | gnomAD v2.1 | Peak Curr. Density (% of WT) | ACMG Classification Pre | ACMG Classification Post |
|---|---|---|---|---|---|---|---|---|
| Wildtype | WT | * | * | * | * | 100 (3.7) | * | * |
| Previously Studied | G9V | 0 | 1 | 0 | 0 | 115.6 (24.7) | VUS | VUS |
| | R121W | 3 | 0 | 0 | 0 | 0.7 (0.6) | LP | LP |
| | T220I | 2 | 0 | 2 | 197 | 86.7 (8.6) | VUS | VUS |
| | T353I | 5 | 0 | 0 | 0 | 0.1 (0.1) | P | P |
| | R620H | 1 | 0 | 3 | 7 | 113 (28.6) | B | B |
| | G752R | 8 | 0 | 1 | 1 | 23.2 (7.1) | P | P |
| | N1325S | 0 | 23 | 0 | 0 | 114.3 (22.4) | P(LQT) | P(LQT) |
| | R1432G | 1 | 0 | 0 | 0 | 2.2 (1) | LP | LP |
| | R1632H | 10 | 0 | 3 | 2 | 64.8 (11.1) | P | P |
| | E1784K | 31 | 114 | 19 | 0 | 51.9 (18.8) | P(LQT +BrS) | P(LQT +BrS) |
| Suspected Benign | A166T | 0 | 0 | 0 | 70 | 103.2 (21.8) | VUS | B |
| | A286S | 0 | 0 | 1 | 79 | 105.8 (12.7) | LB | B |
| | L299M | 0 | 0 | 1 | 59 | 104.3 (16.3) | LB | B |
| | V714A | 0 | 0 | 0 | 16 | 101 (8.9) | VUS | B |
| | G833R | 0 | 0 | 0 | 37 | 109.1 (14.8) | VUS | B |
| | V924I | 0 | 0 | 2 | 31 | 94.8 (12.6) | VUS | LB |
| | V1251M | 0 | 0 | 1 | 59 | 120.5 (13.6) | VUS | B |
| | V1279I | 0 | 0 | 2 | 27 | 103.2 (12.7) | VUS | B |
| | D1370G | 0 | 0 | 0 | 15 | 85.1 (10.6) | VUS | B |
| | A1746V | 0 | 0 | 0 | 15 | 89.4 (12.7) | VUS | B |
| Suspected Brugada Syndrome-associated | D84N | 2 | 0 | 0 | 0 | 32.8 (5) | VUS | LP |
| | F93S | 2 | 0 | 0 | 0 | 0.2 (0.2) | VUS | LP |
| | N109K | 3 | 0 | 0 | 1 | 119.6 (19.5) | VUS | LB |
| | L136P | 2 | 0 | 0 | 0 | 39 (6.4) | VUS | LP |
| | K175N | 1 | 0 | 0 | 0 | 117.8 (14.5) | VUS | VUS |
| | A178G | 1 | 0 | 0 | 0 | 109.6 (19.8) | VUS | VUS |
| | V223L | 2 | 0 | 0 | 0 | 34.2 (6.7) | VUS | LP |
| | L276Q | 2 | 0 | 0 | 0 | 0.8 (0.6) | VUS | LP |
| | R282C | 2 | 0 | 1 | 0 | 1.3 (0.3) | LP | P |
| | C335R | 2 | 0 | 0 | 0 | 0 (0) | VUS | LP |
| | E346D | 1 | 0 | 0 | 0 | 113.2 (13.9) | VUS | LB |
| | D349N | 4 | 0 | 1 | 4 | 28.5 (7.6) | VUS | LP |
| | R367L | 1 | 0 | 0 | 0 | 0 (0) | LP | P |
| | R367C | 3 | 2 | 0 | 3 | 0.6 (0.4) | LP | P |
| | M369K | 3 | 0 | 0 | 0 | 3.4 (0.8) | VUS | LP |
| | G386R | 2 | 0 | 0 | 0 | 1.2 (0.7) | VUS | LP |
| | V396L | 1 | 0 | 0 | 0 | 32 (5) | VUS | LP |
| | M734V | 1 | 0 | 0 | 0 | 68.1 (8.7) | VUS | VUS |
| | A735E | 2 | 0 | 0 | 0 | 0.9 (0.6) | VUS | LP |
| | A735T | 3 | 0 | 1 | 0 | 63.8 (10.1) | VUS | VUS |
| | E746K | 4 | 0 | 0 | 6 | 41.7 (10.8) | VUS | LP |
| | M764K | 2 | 0 | 0 | 0 | 77.5 (8.6) | VUS | VUS |
| | D772N | 1 | 1 | 0 | 5 | 105.3 (10.8) | VUS | VUS |
| | P773S | 1 | 0 | 0 | 0 | 120.5 (10.5) | VUS | VUS |
| | D785N | 2 | 0 | 0 | 0 | 38.9 (7.2) | VUS | LP |

| | | | | | | | | |
|---|---|---|---|---|---|---|---|---|
| | R808C | 3 | 0 | 0 | 2 | 21 (5.1) | VUS | LP |
| | R814Q | 5 | 2 | 8 | 7 | 117.2 (11.7) | LP (BrS) | LP (LQT) |
| | L839P | 4 | 0 | 1 | 0 | 2.9 (2.1) | VUS | LP |
| | F851L | 2 | 0 | 0 | 2 | 16 (2.3) | VUS | LP |
| | W879R | 2 | 0 | 0 | 0 | 0 (0) | VUS | LP |
| | F892I | 2 | 0 | 0 | 0 | 0.8 (0.6) | VUS | LP |
| | E901K | 6 | 0 | 0 | 0 | 3.2 (0.5) | LP | P |
| | N927S | 4 | 0 | 0 | 0 | 29.8 (5.7) | VUS | LP |
| | L928P | 1 | 0 | 0 | 0 | 1.1 (0.8) | VUS | LP |
| | P1014S | 2 | 0 | 0 | 0 | 121.4 (13.2) | VUS | LB |
| | E1225K | 11 | 1 | 1 | 1 | 36 (5.9) | LP | P |
| | D1243N | 5 | 0 | 1 | 41 | 114.7 (15.2) | VUS | B |
| Suspected Brugada Syndrome-associated | G1262S | 4 | 0 | 0 | 8 | 47 (15.5) | VUS | LP |
| | V1281F | 2 | 0 | 0 | 0 | 102.4 (15.5) | VUS | VUS |
| | W1345C | 3 | 0 | 0 | 0 | 12 (2.2) | VUS | LP |
| | L1346P | 2 | 0 | 0 | 0 | 1.6 (0.7) | VUS | LP |
| | V1353M | 2 | 0 | 1 | 7 | 102.6 (16.5) | VUS | B |
| | N1380K | 2 | 0 | 0 | 0 | 0.2 (0.2) | VUS | LP |
| | S1382I | 3 | 0 | 5 | 0 | 3.5 (0.8) | VUS | LP |
| | V1405L | 2 | 0 | 0 | 0 | 13.9 (2.8) | VUS | LP |
| | V1405M | 3 | 0 | 0 | 0 | 36 (6) | VUS | LP |
| | G1406E | 2 | 0 | 0 | 0 | 32.6 (6.2) | LP | P |
| | G1420V | 2 | 0 | 0 | 0 | 0 (0) | VUS | LP |
| | G1420R | 3 | 0 | 0 | 0 | 3 (1.5) | VUS | LP |
| | A1428V | 4 | 0 | 2 | 0 | 0.3 (0.3) | LP | P |
| | Y1449C | 5 | 0 | 4 | 0 | 10.2 (3.4) | LP | P |
| | T1461S | 2 | 0 | 2 | 0 | 59.7 (6.3) | VUS | VUS |
| | E1574K | 4 | 0 | 0 | 0 | 43.3 (12.2) | VUS | LP |
| | R1583C | 2 | 0 | 0 | 2 | 78.9 (7.2) | VUS | VUS |
| | G1642E | 1 | 0 | 0 | 0 | 14.8 (2.5) | VUS | LP |
| | G1661R | 7 | 0 | 0 | 0 | 5.4 (1.5) | LP | P |
| | S1672Y | 3 | 0 | 0 | 0 | 0.9 (0.5) | VUS | LP |
| | A1680T | 4 | 1 | 0 | 13 | 89.5 (14.6) | VUS | B |
| | T1709M | 2 | 1 | 0 | 1 | 23.1 (3.2) | VUS | LP |
| | N1722D | 3 | 0 | 5 | 0 | 37.2 (3.8) | VUS | LP |
| | P1730H | 2 | 0 | 3 | 0 | 45.3 (5.1) | VUS | LP |
| | R1898C | 2 | 0 | 1 | 10 | 28.4 (8.6) | VUS | LP |
| | R1958X | 0 | 2 | 0 | 13 | 59.3 (8.3) | LP | LP |

Peak current density is normalized to WT and presented as mean (standard error).

VUS=Variant of Uncertain Significance, P=Pathogenic, LP=Likely Pathogenic, B=Benign, LB=Likely Benign. Susp. BrS = Suspected BrS, Susp. Benign = Suspected Benign, Unaff.=Unaffected, Peak curr. Density=Peak Current Density. Patient counts are from a literature curation of *SCN5A* variants.[14, 15] Variant classifications are for Brugada Syndrome unless otherwise indicated.

**Table S2: Zone boundaries and restriction enzymes**

| Zone | Size (bp) | Location (bp) | Restriction enzymes |
|---|---|---|---|
| 1 | 727 | 1-727 | AgeI, BssHII |
| 2 | 835 | 722-1557 | BssHII, SpeI |
| 3 | 683 | 1552-2235 | SpeI, EcoRI |
| 4 | 757 | 2230-2987 | EcoRI, NheI |
| 5 | 829 | 2982-3811 | NheI, NsiI |
| 6 | 790 | 3806-4596 | NsiI, AatII |
| 7 | 769 | 4591-5360 | AatII, AflII |
| 8 | 704 | 5355-6059 | AflII, NdeI |

Zone locations are slightly overlapping because each restriction enzyme site is present in two adjacent zone plasmids. Coordinates refer to the canonical numbering scheme with Q1077 included (ENST00000333535), although the wildtype plasmid used in this study had Q1077 deleted (ENST00000443581).

**Table S3 Primers used in this study.**

| Variant | Zone | Name | Sequence |
|---|---|---|---|
| G9V | 1 | ag654 | CCTATTACCTCGGGTCACCAGCAGCTTCC |
| D84N | 1 | ag737 | CCCTGGAGGACCTGAACCCCTTCTATAGC |
| F93S | 1 | ag738 | CTATAGCACCCAAAAGACTTCCATCGTACTGAATAAAGGCA |
| N109K | 1 | ag739 | TTCCGGTTCAGTGCCACCAAAGCCTTGTATGTC |
| R121W | 1 | ag655 | CTTCCACCCCATCTGGAGAGCGGCTGT |
| L136P | 1 | ag740 | CTCGCTCTTCAACATGCCCATCATGTGCACCATCC |
| A166T | 1 | ag656 | TCGAGTACACCTTCACCACCATTTACACCTTTGAG |
| K175N | 1 | ag775 | TGAGTCTCTGGTCAACATTCTGGCTCGAGGC |
| A178G | 1 | ag776 | GGTCAAGATTCTGGGTCGAGGCTTCTGCC |
| T220I | 1 | ag728 | CTCAGCCTTACGCATCTTCCGAGTCCTCC |
| V223L | 1 | ag741 | CCTTACGCACCTTCCGACTCCTCCGGG |
| L276Q | 2 | ag742 | CTCTTCATGGGCAACCAAAGGCACAAGTGCGTG |
| R282C | 2 | ag729 | GGCACAAGTGCGTGTGCAACTTCACAGCG |
| A286S | 2 | ag660 | GTGCGTGCGCAACTTCACATCGCTCAACGG |
| L299M | 2 | ag661 | GGAGGCCGACGGCATGGTCTGGGAATC |
| C335R | 2 | ag785 | GACGCTGGGACACGTCCGGAGGGCT |
| E346D | 2 | ag777 | TAAAGGCAGGCGACAACCCCGACCACG |
| D349N | 2 | ag786 | AGGCGAGAACCCCAACCACGGCTACAC |
| T353I | 2 | ag664 | CCGACCACGGCTACATCAGCTTCGATTCCTT |
| R367L | 2 | ag778 | TTTCTTGCACTCTTCCTCCTGATGACGCAGGAC |
| R367C | 2 | ag665 | CTTTCTTGCACTCTTCTGCCTGATGACGCAGGA |
| M369K | 2 | ag743 | CTCTTCCGCCTGAAGACGCAGGACTGC |
| G386R | 2 | ag745 | AGACCCTCAGGTCCGCAAGGAAGATCTACATG |
| V396L | 2 | ag779 | CATGATCTTCTTCATGCTTCTCATCTTCCTGGGGTC |
| R620H | 3 | ag128 | AAGCCACCTCCTCCACCCTGTGATGCTAG |
| V714A | 3 | ag691 | GGAGTGAAGTTGGTGGCCATGGACCCGTTTACT |
| M734V | 3 | ag780 | CTCAACACACTCTTCGTGGCGCTGGAGCACT |
| A735E | 3 | ag746 | CAACACACTCTTCATGGAGCTGGAGCACTACAACA |
| A735T | 3 | ag747 | TCAACACACTCTTCATGACGCTGGAGCACTACAAC |
| E746K | 4 | ag669 | GCGGCCGCGAATTCAAGGAGATGCTGCA |
| G752R | 4 | ag145 | GGAGATGCTGCAGGTCAGAAACCTGGTCTTCAC |
| M764K | 4 | ag790 | GGATTTTCACAGCAGAGAAGACCTTCAAGATCATTGC |
| D772N | 4 | ag670 | TCAAGATCATTGCCCTCAACCCCTACTACTACTTC |
| P773S | 4 | ag781 | AGATCATTGCCCTCGACTCCTACTACTACTTCCAA |
| D785N | 4 | ag748 | AGGGCTGGAACATCTTCAACAGCATCATCGTCATC |
| R808C | 4 | ag672 | CTTGTCGGTGCTGTGCTCCTTCCGCCT |
| R814Q | 4 | ag674 | CTTCCGCCTGCTGCAGGTCTTCAAGCTGG |
| G833R | 4 | ag675 | ACTCATCAAGATCATCAGGAACTCAGTGGGGGC |
| L839P | 4 | ag749 | CAGTGGGGGCACCGGGGAACCTGAC |
| F851L | 4 | ag799 | TGCTTGCCATCATCGTGCTCATCTTTGCTGTGGTG |
| W879R | 4 | ag798 | CCTGCTGCCTCGCAGGCACATGATGGA |
| F892I | 4 | ag750 | GCCTTCCTCATCATCATCCGCATCCTCTGTG |
| E901K | 4 | ag678 | CTGTGGAGAGTGGATCAAGACCATGTGGGACTG |
| V924I | 4 | ag680 | TGGTCTTCTTGCTTGTTATGATCATTGGCAACCTTGTGGTC |
| N927S | 4 | ag751 | CTTGTTATGGTCATTGGCAGCCTTGTGGTCCTGAAT |
| L928P | 4 | ag782 | TATGGTCATTGGCAACCCTGTGGTCCTGAATCTCT |
| P1014S | 5 | ag792 | CCCCGCCACCCTCAGAGACGGAG |
| E1225K | 5 | ag687 | GGAGCGCTGGCCTTCAAGGACATCTACCTAG |
| D1243N | 5 | ag688 | GGTTCTGCTTGAGTATGCCAACAAGATGTTCACATATGT |
| V1251M | 5 | ag689 | TGTTCACATATGTCTTCATGCTGGAGATGCTGCTC |
| G1262S | 5 | ag690 | TCAAGTGGGTGGCCTACAGCTTCAAGAAGTACTTC |

| | | | |
|---|---|---|---|
| V1279I | 6 | ag704 | GCTCGACTTCCTCATCATAGATGTCTCTCTGGT |
| V1281F | 6 | ag752 | CTTCCTCATCGTAGACTTCTCTCTGGTCAGCCT |
| N1325S | 6 | ag132 | GGCATGAGGGTGGTGGTCAGTGCCCTGGTG |
| W1345C | 6 | ag753 | GTCTGCCTCATCTTCTGCCTCATCTTCAGCATCAT |
| L1346P | 6 | ag754 | CTGCCTCATCTTCTGGCCCATCTTCAGCATCATGG |
| V1353M | 6 | ag697 | CTTCAGCATCATGGGCATGAACCTCTTTGCGGG |
| D1370G | 6 | ag698 | CAACCAGACAGAGGGAGGCTTGCCTTTGAACTACA |
| N1380K | 6 | ag755 | TTTGAACTACACCATCGTGAACAAAAAGAGCCAGTGTG |
| S1382I | 6 | ag795 | CTACACCATCGTGAACAACAAGATCCAGTGTGAGTC |
| V1405L | 6 | ag756 | AGTCAACTTTGACAACCTGGGGGCCGGGTA |
| V1405M | 6 | ag757 | AAAGTCAACTTTGACAACATGGGGGCCGGGTAC |
| G1406E | 6 | ag758 | CTTTGACAACGTGGAGGCCGGGTACCTGG |
| G1420R | 6 | ag759 | GTCCATCCAGCGTTTAAATGTTGCCACCTGCAG |
| G1420V | 6 | ag760 | GCAGGTGGCAACATTTAAAGTCTGGATGGACATTATGTATG |
| A1428V | 6 | ag761 | GGACATTATGTATGCAGTTGTGGACTCCAGGGGG |
| R1432G | 6 | ag783 | CAGCTGTGGACTCCGGGGGGTATGAAGAG |
| Y1449C | 6 | ag762 | ACCTCTACATGTACATCTGTTTTGTCATTTTCATCAT |
| T1461S | 6 | ag796 | CATCTTTGGGTCTTTCTTCTCCCTGAACCTCTTTATTGG |
| E1574K | 7 | ag764 | TGGCCATCTTCACAGGCAAGTGTATTGTCAAGCTG |
| R1583C | 7 | ag794 | AGCTGGCTGCCCTGTGCCACTACTACTTC |
| R1632H | 7 | ag642 | GGCCCGAATAGGCCACATCCTCAGACTGA |
| G1642E | 7 | ag784 | GAGGGGCCAAGGAGATCCGCACGCT |
| G1661R | 7 | ag645 | GCCCTCTTCAACATCAGGCTGCTGCTCTTCC |
| S1672Y | 7 | ag766 | CGTCATGTTCATCTACTACATCTTTGGCATGGCCA |
| A1680T | 7 | ag646 | TTGGCATGGCCAACTTCACTTATGTCAAGTGGGAG |
| T1709M | 7 | ag797 | TTCCAGATCACCATGTCGGCCGGCTGG |
| N1722D | 7 | ag791 | AGCCCCATCCTCGACACTGGGCCGC |
| P1730H | 7 | ag793 | CCCTACTGCGACCACACTCTGCCCAAC |
| A1746V | 7 | ag650 | CGGGAGCCCAGTCGTGGGCATCC |
| E1784K | 7 | ag653 | GGAGGAGAGCACCAAGCCCTTAAGGCG |
| R1898C | 8 | ag700 | CCACCACACTCCGGTGCAAGCACGAAGAG |
| R1958X | 8 | ag734 | GTGAGAACTTCTCCTGACCCCTTGGCCCA |

**Table S4: Summary of the Rosetta energy functions used for $\Delta\Delta G$ calculation**

| Energy term | Weight | Description |
|---|---|---|
| fa_atr | 0.18 | Lennard-Jones attractive between atoms in different residues |
| fa_dun | 0.07 | Internal energy of sidechain rotamers as derived from Dunbrack's statistics |
| fa_mbenv | 0.17 | Statistics-based depth-dependent membrane environment potential |
| fa_mpsolv | 0.23 | Statistics-based depth- and burial-dependent solvation potential |
| fa_pair | 0.57 | Statistics-based amino-acid pair potential, favors salt bridges |
| fa_rep | 0.08 | Lennard-Jones repulsive between atoms in different residues |
| hbond_bb_sc | 0.45 | Sidechain-backbone hydrogen bond energy |
| hbond_sc | 0.43 | Sidechain-sidechain hydrogen bond energy |
| omega | 0.09 | Omega dihedral in the backbone. A harmonic constraint on planarity with standard deviation of ~6 degrees |

### Table S5. All measured parameters for each variant

| Variant | n | Peak density | V1/2 Activation | Inact. time | V1/2 Inactivation | Rec. from Inactivation | Late (% peak) |
|---|---|---|---|---|---|---|---|
| WT | 471 | 100 (3.7) | -36.8 (0.6) | 2 (0) | -95.5 (0.4) | 7.8 (0.3) | 0.53 (0.04) |
| G9V (Prev) | 19 | 115.6 (24.7) | -40.2 (1.1) | 1.8 (0.1) | -100.6 (1.3) | 10.5 (0.8) | 0.46 (0.14) |
| R121W (Prev) | 17 | 0.7 (0.6) | * | * | * | * | * |
| T220I (Prev) | 28 | 86.7 (8.6) | -39.5 (2) | 2 (0.1) | -99.2 (1) | 11.4 (0.7) | 0.4 (0.06) |
| T353I (Prev) | 19 | 0.1 (0.1) | * | * | * | * | * |
| R620H (Prev) | 17 | 113 (28.6) | -44.1 (1.3) | 1.9 (0.2) | -98.2 (1.4) | 9.4 (0.4) | 0.54 (0.1) |
| G752R (Prev) | 14 | 23.2 (7.1) | -14.2 (2.4) | * | * | * | * |
| N1325S (Prev) | 16 | 114.3 (22.4) | -46.4 (1.4) | 2.2 (0.1) | -98.7 (1.9) | 10.2 (0.9) | 1.89 (0.09) |
| R1432G (Prev) | 16 | 2.2 (1) | * | * | * | * | * |
| R1632H (Prev) | 31 | 64.8 (11.1) | -47.7 (2.1) | 2.7 (0.2) | -107.5 (3.9) | 133.9 (13) | 0.44 (0.07) |
| E1784K (Prev) | 12 | 51.9 (18.8) | * | 1.4 (0.2) | -100.9 (3.7) | 6.9 (0.5) | * |
| A166T (S.Ben) | 37 | 103.2 (21.8) | -32.6 (1.9) | 2.7 (0.3) | -104 (1.9) | 7.7 (0.9) | * |
| A286S (S.Ben) | 36 | 105.8 (12.7) | -33.6 (1.4) | 2.3 (0.2) | -92.8 (0.9) | 5.5 (0.8) | 0.83 (0.2) |
| L299M (S.Ben) | 30 | 104.3 (16.3) | -31.6 (1.8) | 3 (0.3) | -92.9 (1.2) | 5.7 (0.6) | 0.73 (0.16) |
| V714A (S.Ben) | 41 | 101 (8.9) | -30.6 (1.2) | 2.5 (0.1) | -90.6 (1.2) | 4.3 (0.8) | 0.51 (0.22) |
| G833R (S.Ben) | 26 | 109.1 (14.8) | -35.5 (1.9) | 2.2 (0.2) | -93.3 (1) | 7.3 (1.5) | 0.59 (0.22) |
| V924I (S.Ben) | 33 | 94.8 (12.6) | -31.6 (1.7) | 2.2 (0.2) | -90.8 (1) | 5.4 (0.6) | 0.89 (0.22) |
| V1251M (S.Ben) | 33 | 120.5 (13.6) | -31.9 (1.2) | 2.5 (0.1) | -91.1 (1.2) | 3.5 (0.2) | 0.58 (0.1) |
| V1279I (S.Ben) | 33 | 103.2 (12.7) | -30.9 (1.7) | 2.5 (0.2) | -92.8 (1) | 5 (0.6) | 0.45 (0.07) |
| D1370G (S.Ben) | 35 | 85.1 (10.6) | -30.6 (1.4) | 2.5 (0.2) | -94.7 (1.5) | 6.5 (0.9) | * |
| A1746V (S.Ben) | 26 | 89.4 (12.7) | -35.4 (2.6) | 2.5 (0.1) | -92.2 (1.3) | 8.6 (1.2) | * |
| D84N (S.BrS) | 16 | 32.8 (5) | -28.9 (3.1) | 3.1 (1.2) | * | * | * |
| F93S (S.BrS) | 15 | 0.2 (0.2) | * | * | * | * | * |
| N109K (S.BrS) | 22 | 119.6 (19.5) | -38.2 (2.6) | 2.5 (0.2) | -94.2 (1.5) | 5.7 (1) | 0.98 (0.1) |
| L136P (S.BrS) | 16 | 39 (6.4) | * | 3.3 (0.7) | * | * | * |
| K175N (S.BrS) | 15 | 117.8 (14.5) | -29.5 (3.8) | 2.2 (0.2) | -104.6 (2.1) | * | * |
| A178G (S.BrS) | 11 | 109.6 (19.8) | -27.6 (1.6) | 2.2 (0.3) | * | * | * |
| V223L (S.BrS) | 14 | 34.2 (6.7) | -39.1 (2.9) | 2.1 (0.1) | -98.5 (3) | * | * |
| L276Q (S.BrS) | 14 | 0.8 (0.6) | * | * | * | * | * |
| R282C (S.BrS) | 67 | 1.3 (0.3) | * | 2.6 (0.6) | * | * | * |
| C335R (S.BrS) | 24 | 0 (0) | * | * | * | * | * |
| E346D (S.BrS) | 30 | 113.2 (13.9) | -44.4 (3.2) | 1.7 (0.1) | -99.6 (2.4) | 13.2 (2) | 0.42 (0.07) |
| D349N (S.BrS) | 21 | 28.5 (7.6) | -26.5 (3.1) | * | * | * | * |
| R367C (S.BrS) | 25 | 0.6 (0.4) | * | * | * | * | * |
| R367L (S.BrS) | 39 | 0 (0) | * | * | * | * | * |
| M369K (S.BrS) | 22 | 3.4 (0.8) | * | * | * | * | * |
| G386R (S.BrS) | 11 | 1.2 (0.7) | * | * | * | * | * |
| V396L (S.BrS) | 31 | 32 (5) | -33.5 (1.4) | 2.9 (0.1) | -94.9 (1.3) | 5.3 (0.6) | * |
| M734V (S.BrS) | 18 | 68.1 (8.7) | -36.1 (2.1) | 1.6 (0.1) | -103.2 (1.4) | 16.2 (1.8) | 0.5 (0.11) |
| A735E (S.BrS) | 12 | 0.9 (0.6) | * | * | * | * | * |
| A735T (S.BrS) | 25 | 63.8 (10.1) | -31.4 (1.3) | 2.4 (0.2) | -91.6 (0.7) | 4.3 (0.4) | 0.41 (0.08) |
| E746K (S.BrS) | 15 | 41.7 (10.8) | -36.3 (1.9) | 2 (0.1) | -97 (1.4) | 7.8 (1.1) | * |
| M764K (S.BrS) | 30 | 77.5 (8.6) | -32.9 (2.1) | 2.7 (0.2) | -92.5 (1.4) | 5.6 (1) | 0.27 (0.05) |
| D772N (S.BrS) | 41 | 105.3 (10.8) | -43 (2.5) | 1.4 (0.1) | -96.2 (1.5) | 8.6 (0.9) | 0.51 (0.08) |
| P773S (S.BrS) | 41 | 120.5 (10.5) | -35.1 (1.7) | 2.4 (0.1) | -99.3 (1.1) | 12.4 (1.6) | 0.27 (0.05) |
| D785N (S.BrS) | 27 | 38.9 (7.2) | -25 (1.5) | 2.7 (0.3) | -102.6 (1.8) | 15.6 (1.4) | 0.46 (0.15) |
| R808C (S.BrS) | 12 | 21 (5.1) | -35.8 (1.9) | 1.6 (0.1) | -104.1 (1.3) | 15.9 (1.4) | * |
| R814Q (S.BrS) | 36 | 117.2 (11.7) | -38.9 (1.5) | 2.1 (0.1) | -95.7 (1.1) | 7.7 (0.8) | 1.36 (0.18) |
| L839P (S.BrS) | 20 | 2.9 (2.1) | * | * | * | * | * |
| F851L (S.BrS) | 26 | 16 (2.3) | -32.5 (1.5) | 1.8 (0.1) | -95.8 (3.1) | 8.8 (3.4) | * |

| | | | | | | | |
|---|---|---|---|---|---|---|---|
| W879R (S.BrS) | 43 | 0 (0) | * | * | * | * | * |
| F892I (S.BrS) | 23 | 0.8 (0.6) | * | * | * | * | * |
| E901K (S.BrS) | 16 | 3.2 (0.5) | -32.8 (3) | 2.2 (0.4) | * | * | * |
| N927S (S.BrS) | 13 | 29.8 (5.7) | * | * | * | * | * |
| L928P (S.BrS) | 27 | 1.1 (0.8) | * | * | * | * | * |
| P1014S (S.BrS) | 34 | 121.4 (13.2) | -46.6 (1.8) | 1.4 (0.1) | -95.9 (1.6) | 8.1 (1.1) | 0.43 (0.07) |
| E1225K (S.BrS) | 19 | 36 (5.9) | -32.9 (1.3) | 2.4 (0.1) | -84 (1.5) | 4.2 (0.5) | * |
| D1243N (S.BrS) | 42 | 114.7 (15.2) | -31.1 (1.5) | 2.6 (0.1) | -87.7 (1.1) | 3.9 (0.9) | 0.66 (0.16) |
| G1262S (S.BrS) | 10 | 47 (15.5) | -49 (2.7) | 2.3 (0.3) | -97.3 (5.3) | 13 (3.2) | * |
| V1281F (S.BrS) | 39 | 102.4 (15.5) | -39.6 (1.9) | 2.5 (0.1) | -96.8 (1) | 8.4 (1.2) | 0.81 (0.16) |
| W1345C (S.BrS) | 10 | 12 (2.2) | * | * | * | * | * |
| L1346P (S.BrS) | 15 | 1.6 (0.7) | * | * | * | * | * |
| V1353M (S.BrS) | 31 | 102.6 (16.5) | -33 (2.3) | 2.2 (0.2) | -94 (1.6) | 7.4 (0.8) | 0.4 (0.11) |
| N1380K (S.BrS) | 25 | 0.2 (0.2) | * | * | * | * | * |
| S1382I (S.BrS) | 29 | 3.5 (0.8) | -40.3 (4) | 2.7 (0.6) | * | 4.1 (1.1) | * |
| V1405L (S.BrS) | 15 | 13.9 (2.8) | -39.4 (2.3) | 1.4 (0.1) | -97.1 (3.2) | * | * |
| V1405M (S.BrS) | 14 | 36 (6) | -26.8 (2.1) | 2 (0.3) | * | * | * |
| G1406E (S.BrS) | 10 | 32.6 (6.2) | -42.7 (1.6) | 1.9 (0.2) | -98 (2.2) | 6.6 (1.2) | * |
| G1420R (S.BrS) | 16 | 3 (1.5) | * | * | * | * | * |
| G1420V (S.BrS) | 11 | 0 (0) | * | * | * | * | * |
| A1428V (S.BrS) | 24 | 0.3 (0.3) | * | * | * | * | * |
| Y1449C (S.BrS) | 12 | 10.2 (3.4) | * | * | * | * | * |
| T1461S (S.BrS) | 41 | 59.7 (6.3) | -35.7 (1.3) | 1.8 (0.1) | -92.5 (1) | 4.5 (0.4) | 0.63 (0.13) |
| E1574K (S.BrS) | 14 | 43.3 (12.2) | -27 (3.7) | * | * | * | * |
| R1583C (S.BrS) | 38 | 78.9 (7.2) | -39.5 (1.1) | 1.8 (0.1) | -96.8 (0.7) | 4.7 (0.3) | 0.46 (0.06) |
| G1642E (S.BrS) | 27 | 14.8 (2.5) | -33 (1.3) | 2.4 (0.2) | -88.9 (2) | 3.3 (0.5) | * |
| G1661R (S.BrS) | 19 | 5.4 (1.5) | * | 1.6 (0.2) | * | * | * |
| S1672Y (S.BrS) | 18 | 0.9 (0.5) | * | * | * | * | * |
| A1680T (S.BrS) | 29 | 89.5 (14.6) | -42.1 (2.6) | 2.2 (0.1) | -96.7 (1.6) | 6.8 (1.1) | 0.95 (0.2) |
| T1709M (S.BrS) | 33 | 23.1 (3.2) | -35.9 (1.3) | 2.4 (0.1) | -91.8 (1.2) | 3.1 (0.3) | * |
| N1722D (S.BrS) | 26 | 37.2 (3.8) | -29.9 (1.4) | 3.3 (0.2) | -89.5 (0.8) | 4.2 (0.4) | 0.25 (0.13) |
| P1730H (S.BrS) | 31 | 45.3 (5.1) | -30.1 (1.7) | 3.1 (0.2) | -88.2 (1.3) | 4.6 (0.6) | 0.35 (0.09) |
| R1898C (S.BrS) | 13 | 28.4 (8.6) | -44.4 (3.9) | 1.6 (0.1) | -100.7 (1.8) | 11.3 (3.5) | * |
| R1958X (S.BrS) | 30 | 59.3 (8.3) | -30.7 (1.7) | 2.6 (0.1) | -93.7 (1.4) | 4.3 (0.8) | * |

Prev. indicates previously studied, S.Ben indicates Suspected Brugada syndrome-associated, and S.BrS indicates Suspected Benign. This dataset is also available in File S1 in a .csv format. N indicates the total number of cells included for the peak current parameter. The number of analyzed cells for the other parameters are presented in File S1. Only variants with at least 5 qualifying cells were included for non-peak density parameters, so most severe loss of function variants are not included for most parameters.

### Table S6. *SCN5A* missense variants with <10% peak current density

| Variant | Location | Peak current (% of WT) This study | Peak current (% of WT) Previous literature | PMID | BrS1 | LQT3 | Unaff. | gnomAD v2.1 |
|---|---|---|---|---|---|---|---|---|
| F93S | N-terminus | 0.2 | * | * | 2 | 0 | 0 | 0 |
| R104Q | N-terminus | * | 0 | 23805106 | 5 | 0 | 0 | 0 |
| R104W | N-terminus | * | 0 | 22739120 | 2 | 0 | 0 | 1 |
| R121W | N-terminus | 0.7 | 0 | 20395683 | 3 | 0 | 0 | 0 |
| T187I | DI S2-S3 linker | * | 0 | 16325048 | 1 | 0 | 0 | 0 |
| L276Q | DI S5-P linker | 0.8 | * | * | 2 | 0 | 0 | 0 |
| R282C | DI S5-P linker | 1.3 | * | * | 2 | 0 | 1 | 0 |
| R282H | DI S5-P linker | * | 5 | 21840964 | 4 | 0 | 0 | 4 |
| C335R | DI S5-P linker | 0 | * | * | 2 | 0 | 0 | 0 |
| T353I | DI S5-P linker | 0.1 | 8.3 | 17198989 | 5 | 0 | 0 | 0 |
| D356N | DI S5-P linker | * | 0 | 16325048 | 9 | 0 | 0 | 1 |
| R367C | DI S5-P linker | 0.6 | * | * | 3 | 2 | 0 | 3 |
| R367L | DI S5-P linker | 0 | * | * | 1 | 0 | 0 | 0 |
| M369K | DI S5-P linker | 3.4 | * | * | 3 | 0 | 0 | 0 |
| G386R | DI S6 | 1.2 | * | * | 2 | 0 | 0 | 0 |
| A735E | DII S1 | 0.9 | * | * | 2 | 0 | 0 | 0 |
| G752R | DII S2 | 23.2 | 6.3 | 12693506 | 8 | 0 | 1 | 1 |
| L839P | DII S5 | 2.9 | * | * | 4 | 0 | 1 | 0 |
| L846R | DII S5 | * | 0 | 22028457 | 1 | 0 | 0 | 0 |
| R878C | DII S5-P linker | * | 0 | 18616619 | 32 | 0 | 4 | 0 |
| R878H | DII S5-P linker | * | 0 | 25904541 | 5 | 0 | 0 | 0 |
| W879R | DII S5-P linker | 0 | * | * | 2 | 0 | 0 | 0 |
| F892I | DII S5-P linker | 0.8 | * | * | 2 | 0 | 0 | 0 |
| R893H | DII S5-P linker | * | 0 | 25904541 | 6 | 0 | 3 | 1 |
| G897E | DII S5-P linker | * | 0 | 25904541 | 0 | 1 | 0 | 0 |
| E901K | DII S5-P linker | 3.2 | * | * | 6 | 0 | 0 | 0 |
| S910L | DII S6 | * | 0 | 24768612 | 4 | 0 | 0 | 1 |
| L928P | DII S6 | 1.1 | * | * | 1 | 0 | 0 | 0 |
| S1218I | DIII S1 | * | 0 | 23424222 | 3 | 0 | 0 | 0 |
| L1346P | DIII S5 | 1.6 | * | * | 2 | 0 | 0 | 0 |
| N1380K | DIII S5-P linker | 0.2 | * | * | 2 | 0 | 0 | 0 |
| S1382I | DIII S5-P linker | 3.5 | * | * | 3 | 0 | 5 | 0 |
| G1406R | DIII S5-P linker | * | 7.7 | 16632547 | 4 | 0 | 1 | 0 |
| G1408R | DIII S5-P linker | * | 0 | 14523039 | 8 | 1 | 19 | 0 |
| G1420R | DIII S5-P linker | 3 | * | * | 3 | 0 | 0 | 0 |
| G1420V | DIII S5-P linker | 0 | * | * | 2 | 0 | 0 | 0 |
| A1428V | DIII S5-P linker | 0.3 | * | * | 4 | 0 | 2 | 0 |
| D1430N | DIII S5-P linker | * | 0 | 23612926 | 2 | 0 | 0 | 0 |
| R1432G | DIII S5-P linker | 2.2 | 0 | 10727653 | 1 | 0 | 0 | 0 |
| I1660V | DIV S5 | * | 1.5 | 17075016 | 5 | 2 | 2 | 0 |
| G1661R | DIV S5 | 5.4 | * | * | 7 | 0 | 0 | 0 |
| S1672Y | DIV S5 | 0.9 | * | * | 3 | 0 | 0 | 0 |
| G1712C | DIV Pore Helix | * | 0 | 28219873 | 1 | 0 | 0 | 0 |
| G1740R | DIV P-S6 Loop | * | 0 | 15057319 | 2 | 0 | 0 | 0 |
| G1743E | DIV P-S6 Loop | * | 0 | 16945804 | 10 | 0 | 0 | 0 |
| G1743R | DIV P-S6 Loop | * | 0 | 15023552 | 12 | 0 | 6 | 0 |

**File S1. Summary of patch clamp data for each variant (.csv)**

For each variant, peak current density (normalized to wildtype), voltage of ½ activation, inactivation time, voltage of ½ inactivation, recovery from inactivation, and late current (% of peak) are presented. Means, standard errors of the mean, and the number of qualifying cells are presented for each parameter. Some parameters have "normalized" values also included which indicates the difference in the parameter value from wildtype. Only variants with at least 5 qualifying cells are included, so most severe loss of function variants are not included for most parameters. In addition, variants are classified into categories based on their functional properties, and each variant's BS3 and PS3 ACMG criteria for Brugada Syndrome (loss of function) or Long QT syndrome (gain of function) is presented.

Literature Cited

1. Pfahnl AE, Viswanathan PC, Weiss R, Shang LL, Sanyal S, Shusterman V, Kornblit C, London B and Dudley SC, Jr. A sodium channel pore mutation causing Brugada syndrome. *Heart Rhythm*. 2007;4:46-53.
2. Song Y, DiMaio F, Wang RY, Kim D, Miles C, Brunette T, Thompson J and Baker D. High-resolution comparative modeling with RosettaCM. *Structure*. 2013;21:1735-42.
3. Barth P, Wallner B and Baker D. Prediction of membrane protein structures with complex topologies using limited constraints. *Proc Natl Acad Sci U S A*. 2009;106:1409-14.
4. Lomize MA, Pogozheva ID, Joo H, Mosberg HI and Lomize AL. OPM database and PPM web server: resources for positioning of proteins in membranes. *Nucleic Acids Research*. 2012;40:D370-D376.
5. Simons KT, Kooperberg C, Huang E and Baker D. Assembly of protein tertiary structures from fragments with similar local sequences using simulated annealing and Bayesian scoring functions. *J Mol Biol*. 1997;268:209-25.
6. Park H, Bradley P, Greisen P, Liu Y, Mulligan VK, Kim DE, Baker D and DiMaio F. Simultaneous Optimization of Biomolecular Energy Functions on Features from Small Molecules and Macromolecules. *Journal of Chemical Theory and Computation*. 2016;12:6201-6212.
7. Kroncke BM, Duran AM, Mendenhall JL, Meiler J, Blume JD and Sanders CR. Documentation of an Imperative To Improve Methods for Predicting Membrane Protein Stability. *Biochemistry*. 2016;55:5002-9.
8. Roushar FJ, Gruenhagen TC, Penn WD, Li B, Meiler J, Jastrzebska B and Schlebach JP. Contribution of Cotranslational Folding Defects to Membrane Protein Homeostasis. *J Am Chem Soc*. 2019;141:204-215.
9. Casadio R, Vassura M, Tiwari S, Fariselli P and Luigi Martelli P. Correlating disease-related mutations to their effect on protein stability: a large-scale analysis of the human proteome. *Hum Mutat*. 2011;32:1161-70.
10. Alford RF, Leaver-Fay A, Jeliazkov JR, O'Meara MJ, DiMaio FP, Park H, Shapovalov MV, Renfrew PD, Mulligan VK, Kappel K, Labonte JW, Pacella MS, Bonneau R, Bradley P, Dunbrack RL, Jr., Das R, Baker D, Kuhlman B, Kortemme T and Gray JJ. The Rosetta All-Atom Energy Function for Macromolecular Modeling and Design. *J Chem Theory Comput*. 2017;13:3031-3048.
11. Richards S, Aziz N, Bale S, Bick D, Das S, Gastier-Foster J, Grody WW, Hegde M, Lyon E, Spector E, Voelkerding K, Rehm HL and Committee ALQA. Standards and guidelines for the interpretation of sequence variants: a joint consensus recommendation of the American College of Medical Genetics and Genomics and the Association for Molecular Pathology. *Genet Med*. 2015;17:405-24.
12. Potet F, Mabo P, Le Coq G, Probst V, Schott JJ, Airaud F, Guihard G, Daubert JC, Escande D and Le Marec H. Novel brugada SCN5A mutation leading to ST segment elevation in the inferior or the right precordial leads. *J Cardiovasc Electrophysiol*. 2003;14:200-3.
13. Benson DW, Wang DW, Dyment M, Knilans TK, Fish FA, Strieper MJ, Rhodes TH and George AL, Jr. Congenital sick sinus syndrome caused by recessive mutations in the cardiac sodium channel gene (SCN5A). *J Clin Invest*. 2003;112:1019-28.

14. Kroncke BM, Glazer AM, Smith DK, Blume JD and Roden DM. SCN5A (NaV1.5) Variant Functional Perturbation and Clinical Presentation: Variants of a Certain Significance. *Circ Genom Precis Med*. 2018;11:e002095.
15. Kroncke BMS, D.; Glazer, A.; Roden, D.; Blume, J. A Bayesian method using sparse data to estimate penetrance of disease-associated genetic variants. *bioRxiv:* https://wwwbiorxivorg/content/101101/571158v1. 2019.
